# Functional diversification of ERF transcription factors in land plants

**DOI:** 10.64898/2026.10.13.758753

**Authors:** Ignacy Bonter, Akira Iwase, Vanda Adamkova, Kimitsune Ishizaki, Takayuki Kohchi, Keiko Sugimoto, Haseloff J Jim, Facundo Romani

## Abstract

The ERF family is one of the largest families of plant-specific transcription factors, controlling diverse developmental processes and stress responses. Extensive gene duplication and redundancy in flowering plants have generated large ERF repertoires, complicating comparisons of ERF function across land-plant lineages. How ERF transcription factor repertoires and their functions have evolved across land plants remains poorly understood. Here, we reconstruct the phylogenetic relationships and expansion of ERF transcription factors across land plants and exploit the compact ERF repertoire of the model liverwort *Marchantia polymorpha* to investigate their functional diversification. Systematic gain-of-function analysis reveals that different clades of ERFs can drive distinct cell fate and reprogramming outcomes in Marchantia, in addition to stress-related functions. Interestingly, we found that Mp*ERF2*, the sole Group I ERF in Marchantia, phylogenetically related to the WIND clade, promotes rhizoid cell differentiation. This contrast with the role of the WIND clade in wound-induced regeneration in Arabidopsis. Heterologous expression of At*WIND1* in Marchantia was also capable of inducing rhizoid formation, supporting the hypothesis that this reprogramming capacity is conserved at the protein level. On the other hand, loss of Mp*ERF2* reduced rhizoid coverage but did not significantly impair regeneration. These findings provide an evolutionary framework for understanding the functional evolution of ERF transcription factors and how conserved transcriptional regulators are co-opted into novel developmental programs during the radiation of land plants.

## INTRODUCTION

The ETHYLENE RESPONSE FACTOR (ERF) family is the second largest group of transcription factors in land plants (1). Together with AP2 and RAV, ERFs are part of the AP2/ERF superfamily and are characterized by a single AP2/ERF DNA-binding domain. They were originally identified through their association with GCC-box-containing ethylene-responsive genes (2). Although their name reflects this historical connection, ERF proteins function in a much broader range of biological processes, including regeneration, stem cell activity, and responses to a variety of biotic and abiotic stresses, often acting downstream of hormonal pathways such as ethylene, jasmonate, and abscisic acid (3).

ERF genes are subdivided into 10 groups or subfamilies (I-X) (4). The large size and evolutionary diversification of the ERF family have made it challenging to assess how ERF functions have diversified across the family, and most functional studies have therefore focused on individual ERF genes or specific groups. Comparisons based on loss-of-function phenotypes are further complicated by functional redundancy, which can differ among species and between paralogues. In contrast to the 121 ERF genes in *Arabidopsis thaliana*, the liverwort *Marchantia polymorpha* encodes a compact set of only 22 ERF genes (5), with representatives of most groups but reduced within-clade redundancy.

A prominent function of ERF transcription factors is the induction of callus after wounding (6). This has been well described during sporophyte regeneration in flowering plants, and involves ERFs from groups I, VIII, IX, and X. Among them, *ENHANCER OF SHOOT REGENERATION 1 (ESR1*), *WOUND INDUCED DEDIFFERENTIATION 1 (WIND1*), *ERF109*, and *ERF115* (7) are the best characterized examples. This process also involves extensive hormone signalling, cell reprogramming, chromatin remodelling, and the reactivation of stem cell programmes (8, 9). Less is known about regeneration in other clades of land plants, particularly gametophytic-dominant plants such as bryophytes. Recent work in Marchantia has begun to address this question by showing that regeneration recruits ERF transcription factors homologous to known regulators of regeneration in angiosperms. Mp*ERF15*, related to *ERF109* (10), and Mp*ERF20/LOW-AUXIN RESPONSIVE (LAXR)*, an orthologue of *ESR1*, are both induced upon injury and are important for regeneration in Marchantia (11), paralleling the functions of their Arabidopsis counterparts. These findings suggest that components of the ERF regulatory machinery associated with wound-induced regeneration are conserved across deeply divergent land-plant lineages. However, bryophytes such as liverworts and mosses regenerate from the gametophyte with exceptional speed (12). These contrasting regeneration strategies raise a fundamental question about how the underlying gene regulatory networks (GRNs) evolved.

Among the ERF subfamilies most prominently associated with regeneration in angiosperms, group I stands out. In Arabidopsis, this group is represented by *WIND1* and its close paralogues *WIND2–4* are rapidly induced upon wounding and are sufficient, when overexpressed, to trigger ectopic callus formation, promote cellular dedifferentiation, and enhance somatic embryogenesis and shoot regeneration in tissue culture (13–16). WIND1 achieves this by activating cell reprogramming programs, partly through the induction of cytokinin signalling. This positions WIND1 at the junction between wound-induced ERFs and shoot formation via ESR1 (16).

Determining which ERF functions are conserved and which represent lineage-specific innovations is therefore essential for understanding how ERF transcription factors evolve. In Marchantia, there is a single member of the WIND clade, represented by Mp*ERF2*. However, its function is still unknown, as is the case for several other ERFs in this emerging model species. Marchantia therefore provides an opportunity to systematically compare the functional capacities of ERF proteins across the entire family in a deeply divergent land-plant lineage. Gain-of-function analysis provides a particularly useful complementary approach to loss-of-function genetics for comparing ERF proteins across phylogenetic groups, as it tests the activities that individual proteins are sufficient to induce in a common genetic background. This can reveal functional capacities that may be masked by redundancy in loss-of-function analyses and provides a common framework for assessing functional diversification across the family.

Here, we combine phylogenetic reconstruction, genome-wide expression profiling, and systematic gain-of-function analysis across the ERF family to establish an updated evolutionary and functional framework for ERFs in land plants. Our phylogenetic analysis supports an expanded classification of ERFs into 11 groups (I-XI), resolving their relationships and evolutionary history. Analysis in Marchantia revealed extensive functional diversification of ERFs transcription factors, with different subfamilies producing distinct developmental and cellular outcomes, while others retain roles associated with stress and wound-induced regeneration. In addition to Mp*ERF13* and Mp*ERF14/GEMMIFER (*Mp*GMFR)*, which are respectively associated with oil body cell differentiation and gemma formation (17, 18), we found that Mp*ERF2* promotes rhizoid cell differentiation. Remarkably, heterologous overexpression of the Arabidopsis WIND1 also induced ectopic rhizoid formation in Marchantia, showing that both proteins retain the capacity to induce this cell fate activity, despite the deep evolutionary divergence between the two proteins. More broadly, our findings provide an updated view of the functional conservation and diversification of ERF transcription factors in land plants and show how they can be redeployed into distinct developmental programs during the evolution of plant gene regulatory networks.

## RESULTS

### Phylogenetic relationships and diversification of ERF transcription factors

To establish the evolutionary relationships of ERF transcription factors across land plants, we identified ERF genes in two flowering plants (*A. thaliana* and *Amborella trichopoda)*, one fern (*Ceratopteris richardii*), one gymnosperm (*Ginkgo biloba*), one lycophyte (*Selaginella kraussiana*), and four bryophytes (*M. polymorpha, Physcomitrium patens, Anthoceros agrestis,* and *Takakia lepidozioides)*. In total, we classified 628 sequences as part of this family across the species analysed (Figure 1; Supplemental File 1). Our phylogenetic analysis is broadly consistent with the previous classification in 10 groups (I-X) (4), and shows strong support for the different clades. One key difference is the classification of the Arabidopsis RAP2.11 as a separate clade, previously assigned to the single member of group Vb. This group, that we named group XI, is strongly supported by bootstrap and its presence in all species analysed. There is a divergent clade that exist only in mosses, represented in the phylogeny by a single member in Takakia (Figure 1, Supplemental File 1), referred to as “moss-specific” (19).

**Figure 1.**
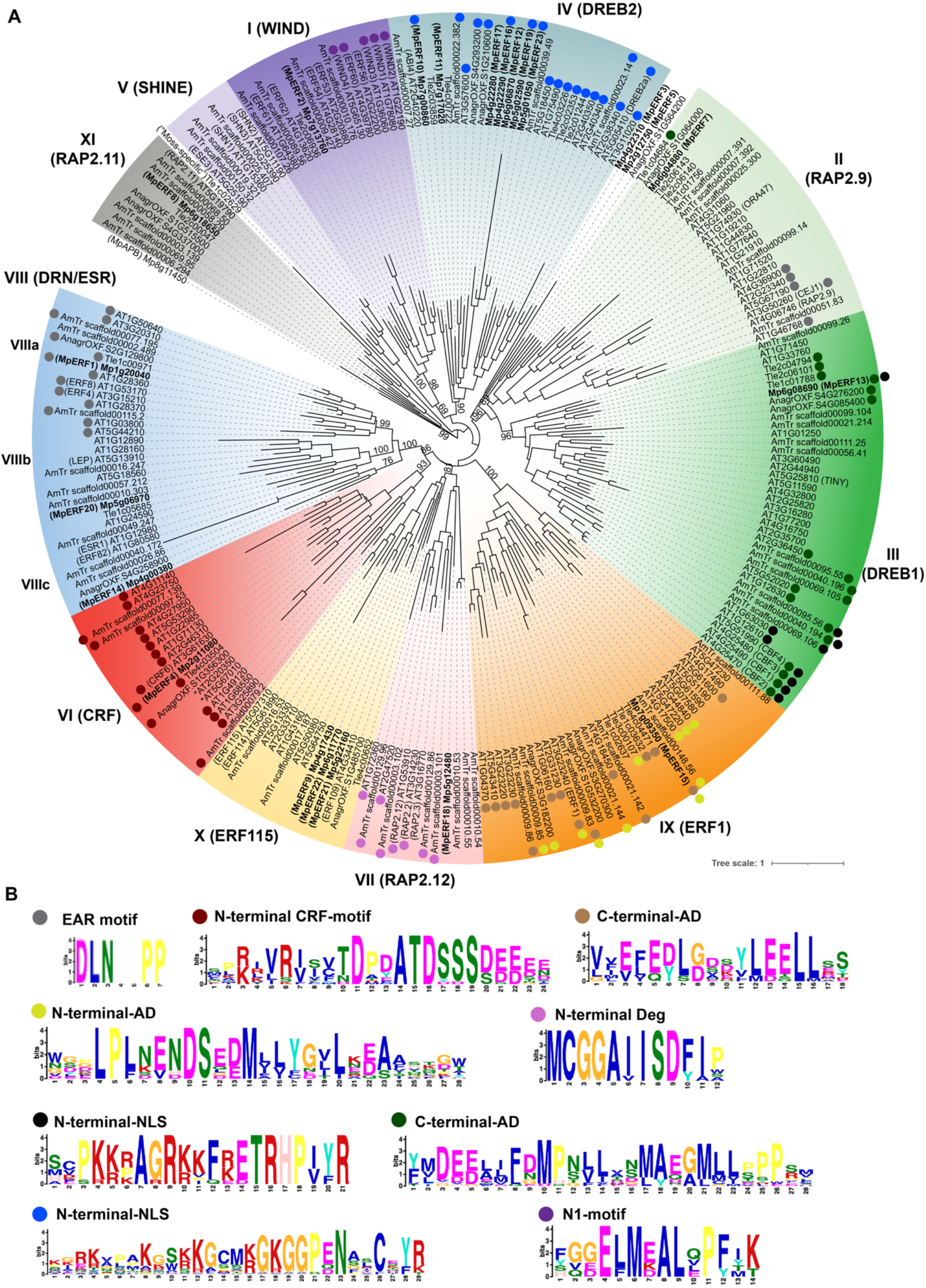
Phylogenetic relationships and expression dynamics of ERF transcription factors in Marchantia. (A) Maximum likelihood phylogeny of ERFs from selected species rooted to MpAPB (AINTEGUMENTA, PLETHORA and BABY BOOM). Gradient colours represent each group as indicated. Relevant bootstrap values are shown. Colored circles indicate the presence of motifs as in (B). (B) Selected logo plots of conserved motifs across ERF groups. AD: Activation domain. NLS: Nuclear localization signal. * Members of the Xb-like group not consistently classified across the larger phylogeny shown in Supplemental File 1. Abbreviations: *Arabidopsis thaliana = AT; Takakia lepidozioides = Tle; Marchantia polymorpha = Mp; Anthoceros agrestis (Oxford strain) = AnagroOXF; Amborella trichopoda = AmTr*.

All conserved 11 groups (I-XI) are present in at least one species of bryophytes, suggesting that this diversification occurred early in the evolution of land plants. ERF genes are subsequently substantially expanded in flowering plants but remain relatively compact in Marchantia. Duplications in groups IV and X in Marchantia are lineage specific and not ancestral to all land plants. On the other hand, group VIII duplications predate the diversification of embryophytes (land plants). We identified three distinct clades (VIIIa, VIIIb, and VIIIc) with members in all bryophyte lineages, rather than the two reported previously (4). Group VIIIc includes MpERF14/GMFR and ERF82 (18); group VIIIb contains ESR1/DRN and its orthologue in Marchantia MpERF20/LAXR (11); and group VIIIa, which includes the Marchantia MpERF1 and, as its closest Arabidopsis homologues, it is characterized by the presence of a repressive EAR motif (DLNxxP) (20). Based on the sampled genomes, our phylogenetic analysis is consistent with at least 13 conserved ERF lineages present in the common ancestor of land plants, corresponding to groups I, II, III, IV, V, VI, VII, VIIIa, VIIIb, VIIIc, IX, X, and XI.

Group IX includes MpERF15, which was previously assigned to group X and considered related to ERF109 (10). Group IX includes several ERFs playing a variety of functions. MpERF15 protein possesses both the conserved C- and N-terminal domains present across the group (Figure 1). ERF1 in particular is one of the best-characterized genes in this clade and is the canonical target of EIN3 in the ethylene signalling pathway and it is also integrated in the jasmonate wound-responsive signalling in Arabidopsis (21, 22). In that sense, our phylogeny instead places MpERF15 as a putative orthologue of ERF1, rather than ERF109. However, regardless of its group assignment, it is consistent with the functional conservation of MpERF15 in Marchantia (10).

In particular, the Marchantia genome encodes for 22 genes, numbered from MpERF1 to MpERF23. MpERF6 represents a deprecated annotation subsequently identified as overlapping with MpERF5 in the revised MpTak1 v5.1 genome annotation (23). ERF groups I, II, III, VI, VII, and IX are represented by a single copy in Marchantia, except for group V which is absent. This group is also absent from 3 out of 4 bryophyte genomes analysed, with *P. patens* as the exception, suggesting it may have been lost repeatedly during the evolution of land plants. Marchantia genome also displays two ERFs, MpERF3 and MpERF5, that we were unable to reliably classify.

### Conserved accessory motifs in ERFs transcription factors

In addition to group VIIIb EAR motif, some members of group II in Arabidopsis are also characterized by the presence of an EAR motif in Arabidopsis (24). However, this motif does not seem to be conserved in the other species analysed here (Figure 1), suggesting that it might have been incorporated later in the diversification of flowering plants.

To extend the analysis to other motifs, we searched for conserved protein motifs outside the AP2/ERF DNA-binding domain using MEME (25). We focused on those whose function can be inferred from the literature and are specific of a given clade. In group III, there are two conserved C-terminal and N-terminal motifs, particularly CBF homologues. It was shown that these motifs act as strong trans-activation domain (26) and nuclear localization signal (NLS) respectively (27). Similar motifs also exist in member of this group in bryophytes, particularly Marchantia harbours both (Figure 1).

In group IV, there is a conserved N-terminal domain that may include the NLS described before for DREB1 TFs (28). This domain is present in both bryophytes and angiosperms but not conserved in the divergent ABI4 (Figure 1).

On the other hand, group VI, also referred to as cytokinin response factors (CRF), contain an N-terminal motif that is characteristic of this clade. This domain is important for protein-protein interactions to form homo and hetero-dimers but also with histidine-phosphotransfer proteins (HPs), key components of the cytokinin two-component signal transduction pathways. The latter interaction is critical for the nuclear translocation of the CRFs (29, 30). Interestingly, this domain is also conserved across land plants (Figure 1).

Group VII ERFs have a conserved N-terminal motif associated with its oxygen-dependent proteasomal degradation (31). However, this motif is not conserved in bryophytes. This is consistent with previous analysis (32).

Group IX ERF proteins could contain acidic domain both in the C-terminal and N-terminal (33). In particular, the C-terminal EDLL motif that acts as a strong trans activator (34). In bryophytes, these acidic domains are also present, either in the C-terminal, the N-terminal, or, as in the case of Marchantia, in both (Figure 1). Interestingly, the Marchantia homologue, possess the same domain architecture as the Arabidopsis ERF1.

Finally, some members of the group I ERFs, particularly WIND1-4, are characterized by an additional N-terminal functional domain denominated N1 (15, 35). The conservation of this motif was not detected by global MEME analysis, but it is enriched among the group I ERFs. Interestingly, N1 is absent from bryophytes proteins such as Marchantia, whereas other conserved motifs are retained (Supplemental Figure 1).

Our motif distributions are consistent with most characteristic features being ancestral, while the group VII degron and group I N1-motif likely arose later. This is consistent with the idea that the minimal set of ERFs in the common ancestor of land plants already contained most of the characteristic protein features associated with the function of each group.

**Supplemental File 1.** Maximum likelihood phylogeny of ERFs from all species tested and rooted to MpAPB (AINTEGUMENTA, PLETHORA and BABY BOOM). Colours represent each group as in Figure 1. Abbreviations: *Arabidopsis thaliana = AT; Takakia lepidozioides = Tle; Marchantia polymorpha = Mp; Anthoceros agrestis (Oxford strain) = AnagroOXF; Amborella trichopoda = AmTr; Physcomitrium patens = Pp; Ceratopteris richardii = Ceric; Selaginella kraussiana = SK; Ginkgo biloba = Gibil*.

### ERFs are dynamically expressed during wounding and regeneration

RNA-seq analysis during regeneration revealed distinct temporal patterns of ERF expression, with some genes responding rapidly after wounding and others increasing at later stages (Figure 1B). As noted previously, several ERFs showed rapid, transient induction within 1-2 h of wounding, including MpERF1/7/8/9/11/15/18/21 (10, 11). These rapidly induced genes span multiple ERF groups (II, VII, IX, X, XI), indicating that the early wound response is not restricted to a single evolutionary lineage. All these genes except for MpERF8 are also significantly induced in plants treated with exogenous dn-OPDA (36), the jasmonate analogue in Marchantia (37), consistent with dn-OPDA acting as a signal contributing to their wound-induced activation. MpERF20 showed a later increase, peaking at 6 h (11, 38). Several of these ERFs have counterparts in Arabidopsis, consistent with conservation of ERF-mediated wound signalling across land plants and the canonical jasmonate/ethylene response (39).

Most lineage-specific ERF duplicates in groups IV and X were either undetectable or weakly expressed during regeneration. Although several are expressed in sporelings and reproductive tissues (Supplemental Figure 2), indicating that their limited expression in this context does not necessarily reflect pseudogenization.

Other ERFs are upregulated at later stages of regeneration, particularly MpERF4, or just fluctuate across the time course. Notably, this group includes MpERF13 and MpERF14/GMFR, which have previously been associated with specific cell identity programs in Marchantia (17, 18). Surprisingly, MpERF2 is among this later group too, with only a modest peak at 6 hours but less than 2-fold compared to time zero. This response is considerably weaker compared with the rapid wound induction of Arabidopsis WIND1 (16), suggesting that the two proteins, despite their phylogenetic relationship, are deployed in distinct developmental contexts. The developmental function of MpERF2 in Marchantia, however, remains unknown. Together, these expression profiles reveal substantial temporal and lineage-specific differences in ERF deployment during regeneration, motivating a systematic functional analysis of ERF activity in Marchantia.

### Inducible overexpression of ERFs reveals their distinct roles in plant development

To systematically explore the developmental functions of the Marchantia ERF family, we generated inducible gain-of-function lines for most ERFs using the _pro_E2F:XVE induction system (11). We cloned a fusion of the CDS corresponding to each ERF with a fluorescent protein and generated stable transgenic lines in Marchantia. Out of 22 Marchantia ERFs, we targeted 18 ERF genes spanning all groups and obtained lines with detectable induction for 15 (Supplemental Figure 3).

Inducible expression produced a broad range of developmental phenotypes after seven days of induction (Figure 3; Supplemental Figure 4), revealing extensive functional diversification across the family. Different ERF genes and groups affected cell differentiation, meristem formation and growth, and related ERFs could produce markedly different developmental outcomes. Conversely, similar phenotypes arose from ERFs belonging to distinct phylogenetic groups.

Among all ERFs tested, MpERF2 (Group I) produced the most distinctive phenotype. Strikingly, overexpression of MpERF2 disrupted normal development and induced widespread ectopic rhizoid-like outgrowths across the epidermis (Figure 3). This phenotype was distinct from the developmental defects produced by other ERFs and suggested that MpERF2 has a specific role in epidermal cell differentiation. This was particularly unexpected given that its closest characterized angiosperm relatives, the WIND proteins, have prominent roles in cellular reprogramming during regeneration (16).

Group VIII members have been previously characterized (11, 18) with strikingly divergent developmental effects. Consistent with the previous report (11), we observed that overexpression of MpERF20/LAXR (group VIIIb) was sufficient to induce ectopic meristems, at least in one of the two independent lines obtained, forming a continuous meristem along the edge of the thallus, (Figure 3). This stem-cell activity was also recently found in *P. patens* (40) and resembles the function of ESR1 in Arabidopsis (41), suggesting it is functionally conserved across land plants. On the other hand, overexpression of MpERF14/GMFR (group VIIIc) caused a distinct developmental phenotype, forming ectopic gemma, as previously shown (18). Overexpression of the remaining member of group VIII, MpERF1 (group VIIIa), showed a strong growth arrest phenotype but not extra meristems (Figure 3). MpERF1 belongs to a subclade characterized by an EAR motif, and its putative Arabidopsis orthologue ERF8 acts as a transcriptional repressor in stress responses (42). This is consistent with the possibility that the distinct phenotype of MpERF1 reflects the repressive regulatory activity associated with this subclade. Similar results were obtained in MpERF1-GR glucocorticoid induced overexpression (18). These phenotypes illustrate how Group VIII ERFs may have undergone substantial functional diversification early during land-plant evolution, resulting in clearly distinct developmental outcomes.

Ectopic meristem formation was not restricted to Group VIII, as MpERF15 (Group IX) also induced ectopic meristems (Figure 3). This is consistent with its established role in regeneration (10). As proposed before, its function might be upstream of MpERF20/LAXR (10, 11), partially resembling the wound induced regeneration network in Arabidopsis (8).

Group III, MpERF13, caused strong growth arrest (Figure 3) but also was able to generate ectopic oil body cells as demonstrated by BODIPY staining (Supplemental Figure 5). This is consistent with T-DNA gain-of-function lines described before (17). These results confirm that ectopic MpERF13 expression is sufficient to induce oil-body differentiation. Oil bodies are a liverwort-specific feature (43) that cannot be directly compared with the biological function of the DREB homologues in Arabidopsis, normally associated with abiotic stresses (44). This illustrates how ERF activity can be deployed into lineage-specific cell differentiation programs, like MpERF14 in gemma formation (18).

Among other members of this family that were not characterized before, none of the other ERFs showed an analogous phenotype in ectopic meristem formation. In particular, MpERF4 and MpERF18 (group VI and VII respectively) are indistinguishable from the wild-type (Figure 3a) and growth is not significantly impaired (Figure 3b). MpERF4 belongs to the CRF family and require interacting with HPs to translocate to the nuclei and be active (29). In the case of MpERF18 (group VII), its homologues in Arabidopsis are associated with the response to hypoxia and tightly controlled post-translationally in an oxygen-dependent manner (31). MpERF18 lacks the conserved N-terminal motif required for its proteasomal degradation by the N-end rule pathway. In Arabidopsis, overexpression of versions of RAP2.12 without the N-terminal motif causes strong growth arrest (3). Yet, we have not seen an analogous phenotype in Marchantia (Figure 3), suggesting that the protein might not be functionally conserved (32).

Among group X, all three members showed a strong growth arrest. Some degree of redundancy is expected as they correspond to liverwort-specific duplications. Yet, in the case of MpERF9 and MpERF22, we observed the accumulation of brown pigments (Figure 3a), likely due to cell death. In the case of MpERF21, we observed a bulge in the apical notch after 7 days in β-estradiol (Figure 3a).

A similar phenotype was also observed for MpERF7 (Figure 3), from group II. Both group II and X are rapidly induced upon wounding and in OPDA and are candidates for conserved components of the jasmonate/ethylene-associated wound-response GRN. This is clearer for group X, where the homologues in Arabidopsis are well described to be involved in the wounding response (39). In the case of group II, some members of the family such as CEJ1 (4) and ORA47 (45), are also associated with the jasmonate/ethylene response.

The sole Group XI member tested, MpERF8, caused strong growth arrest upon induction (Figure 3), but did not reproduce the developmental phenotypes observed for ERFs in other groups. Little is known about this novel group. The single member of this group in Arabidopsis, RAP2.11 (At5g19790), seems to play a role in the response to low potassium (46).

For group IV ERFs in Marchantia, we obtained detectable induction for MpERF10 and MpERF12. Transgenic lines overexpressing MpERF10 MpERF12 showed a reduced thallus size. The induction of MpERF10 also caused other pleiotropic developmental effects, including more rhizoids, but no obvious defects were associated with MpERF12 (Figure 3). In Arabidopsis, group IV ERFs are associated with ABA and stress responses (3), the phenotype of growth retardation could be consistent with those observed in the overexpression of DREB2 (28). On the other hand, MpERF11/17/23 did not show any detectable expression after induction. A knock-out for the Marchantia MpERF23 has been recently described (32). However, no conspicuous phenotypes were reported.

### MpERF2 is expressed in proximal rhizoids during thallus development

Transcriptional reporters of Marchantia ERFs (38) have revealed that most of them are ubiquitously expressed (MpERF1/4/8/21) or around the apical notch (MpERF7/10/11/14/15/17/18/19/20) at the gemma stage. Some are not detected (MpERF3/12/16/23), consistent with RNA-seq analysis (Figure 2). Interestingly, only MpERF13 and MpERF2 are expressed in oil bodies and proximal rhizoids, respectively. Both expression patterns are consistent with their putative functions observed in overexpression analysis (Figure 3).

**Figure 2.**
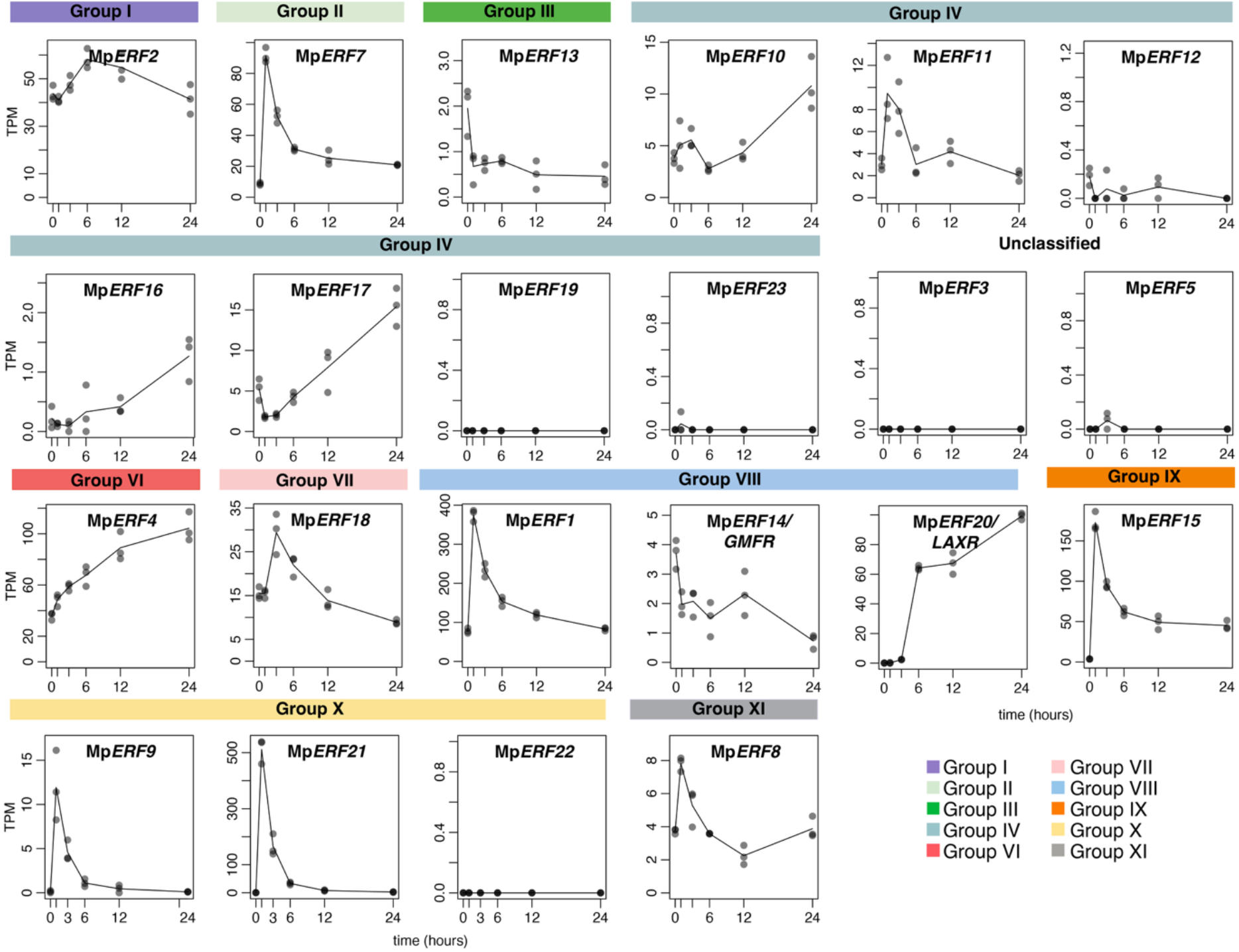
ERF gene expression during regeneration. Expression of *Marchantia polymorpha* ERF genes in the basal region of the thallus at different time points following wounding. Colours represent each group as in Figure 1.

**Figure 3.**
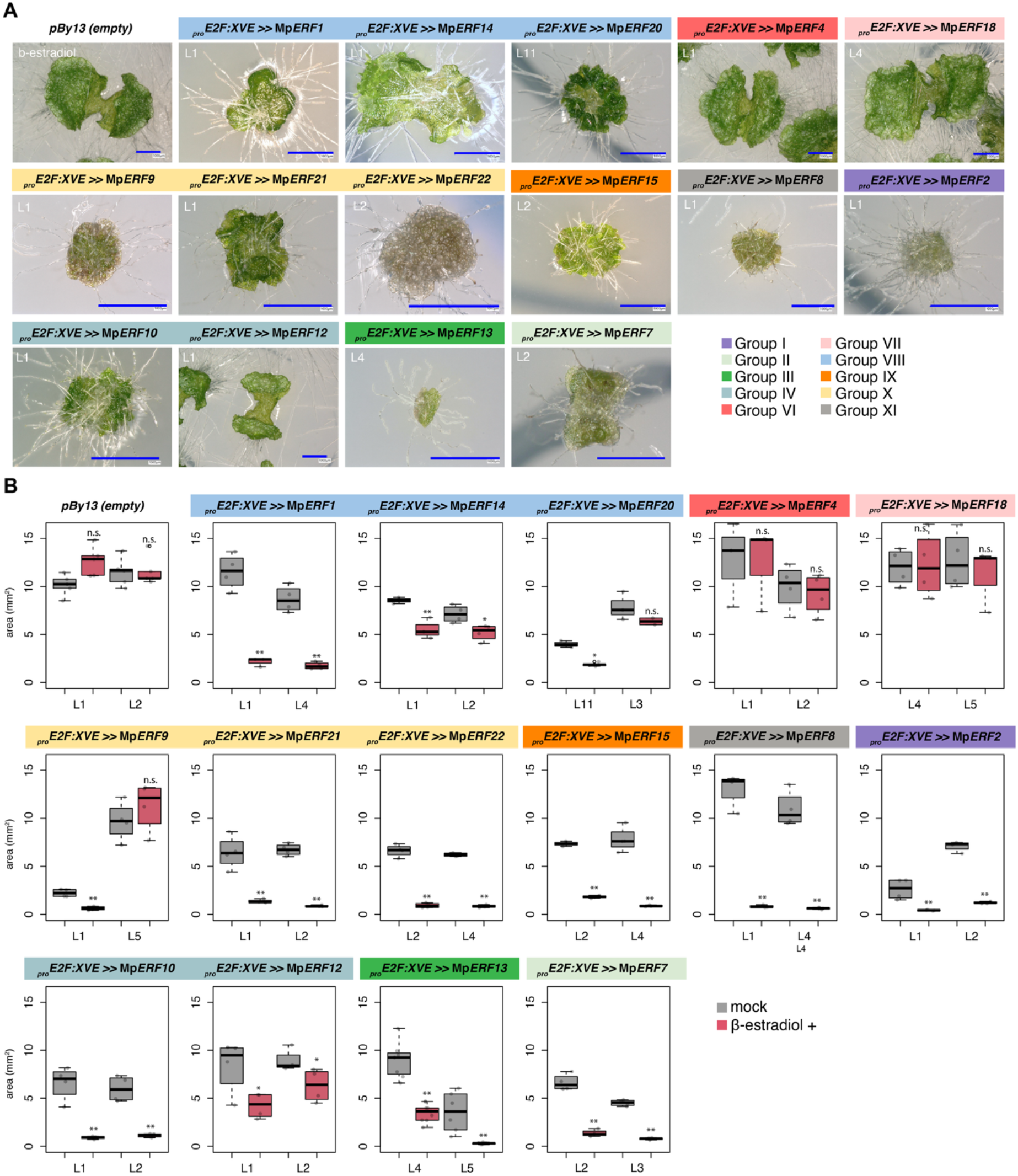
Systematic inducible overexpression of a subset of ERF transcription factors in Marchantia. (A) Representative images of gemmalings of Marchantia ERF inducible lines after 7 days in Gamborg B5 media supplemented with 5 μM β-estradiol. pBy13 correspond to an empty vector control. Colour bars indicate ERFs groups according (see legend). Representative pictures of mock controls and independent transgenic lines are shown in Supplemental Figure 4. (B) Thallus area of the same plants (n = 4). Mock-treated plants are also shown. Two independent lines are shown for each genotype. Colours indicate ERF subfamilies as in panel A. Statistical significance was assessed using Tukey’s HSD test; groups with significantly different means are indicated by different letters (95% confidence level). Scale bars = 100 μm.

At the gemma developmental stage, rhizoids precursors are present on both the dorsal and ventral surfaces as dorso-ventral polarity has not yet been fully established. To gain more insights on MpERF2 expression, we therefore followed *_pro_*Mp*ERF2* (1.8 kb upstream of the ATG, including the 5′ UTR) activity during subsequent development in plants grown either upright or upside down, allowing promoter activity to be monitored on both surfaces. Activity was detected in rhizoid precursors on both surfaces during the first 2 days, but after 3 days was retained predominantly in ventral rhizoid-forming regions as new rhizoids emerged (Figure 4A). At later stages, *_pro_*Mp*ERF2* activity was strongly detected in differentiated rhizoids. Weak *_pro_*Mp*ERF2* activity was also detected in the apical notch but was at least an order of magnitude lower than that observed in rhizoids (Figure 4B). Overall, the spatial expression of MpERF2 in rhizoids is consistent with the idea of a putative role for MpERF2 in rhizoid differentiation in Marchantia. This is also consistent with recent transcriptomic profiling of rhizoid samples (47).

**Figure 4.**
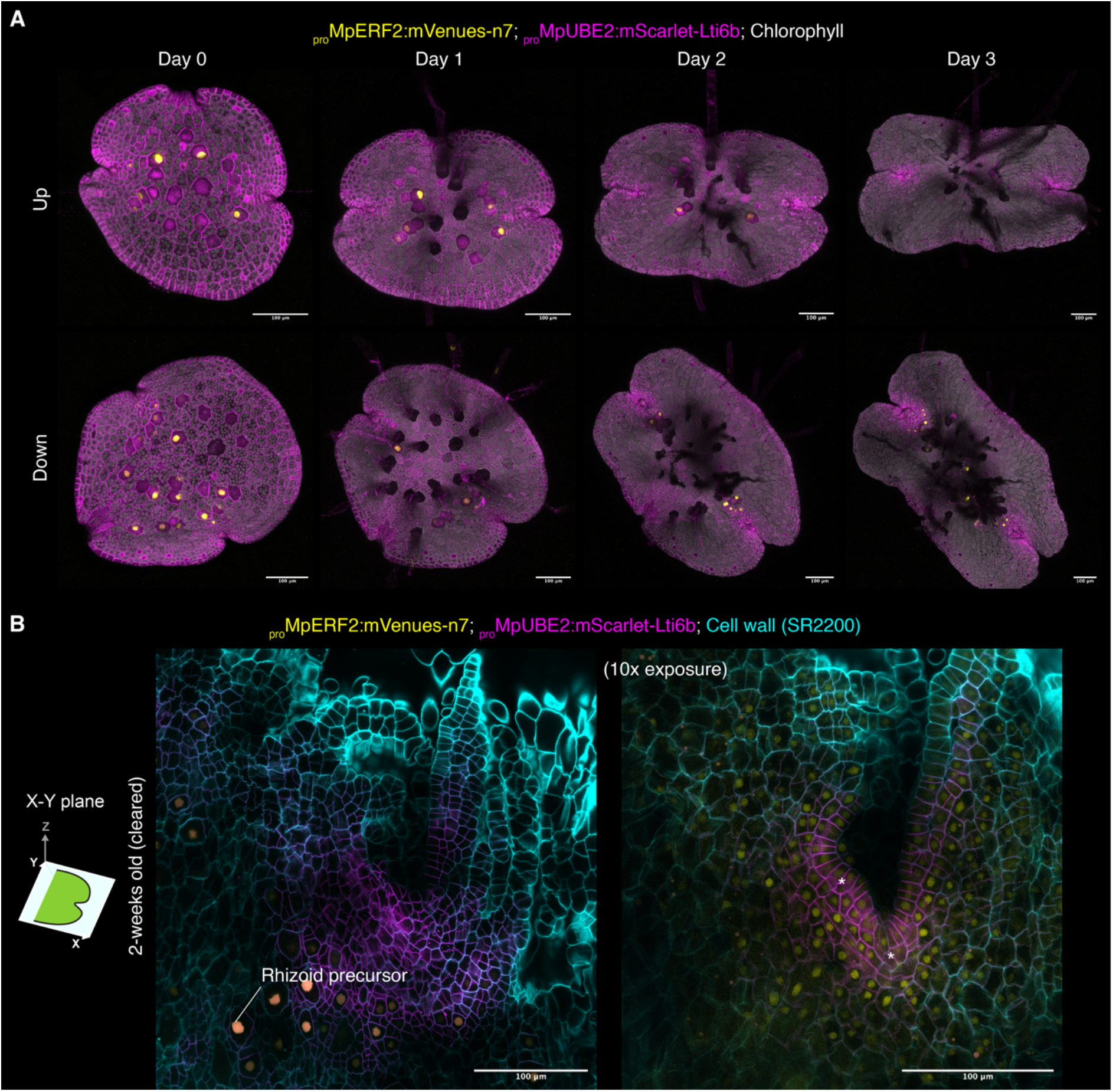
Expression pattern of MpERF2. (A) Maximum-projection confocal images showing *_pro_*Mp*ERF2* activity in dorsal (top) and ventral (bottom) surfaces of gemmalings at different developmental stages. (B) Single optical section of a cleared meristem from a 2-week-old *_pro_*Mp*ERF2* reporter line. * indicate apical cells. Scale bars = 100 μm.

### Heterologous overexpression of AtWIND1 phenocopies MpERF2-induced ectopic rhizoid formation

To further characterize the Mp*ERF2* gain-of-function phenotype, we tested a shorter induction period (3 days) and lower β-estradiol concentrations. Ectopic rhizoid-like outgrowths were observed even at the lowest concentration tested and increased with induction strength, while overall thallus growth was progressively reduced in a dose-dependent manner (Figure 5A,B).

**Figure 5.**
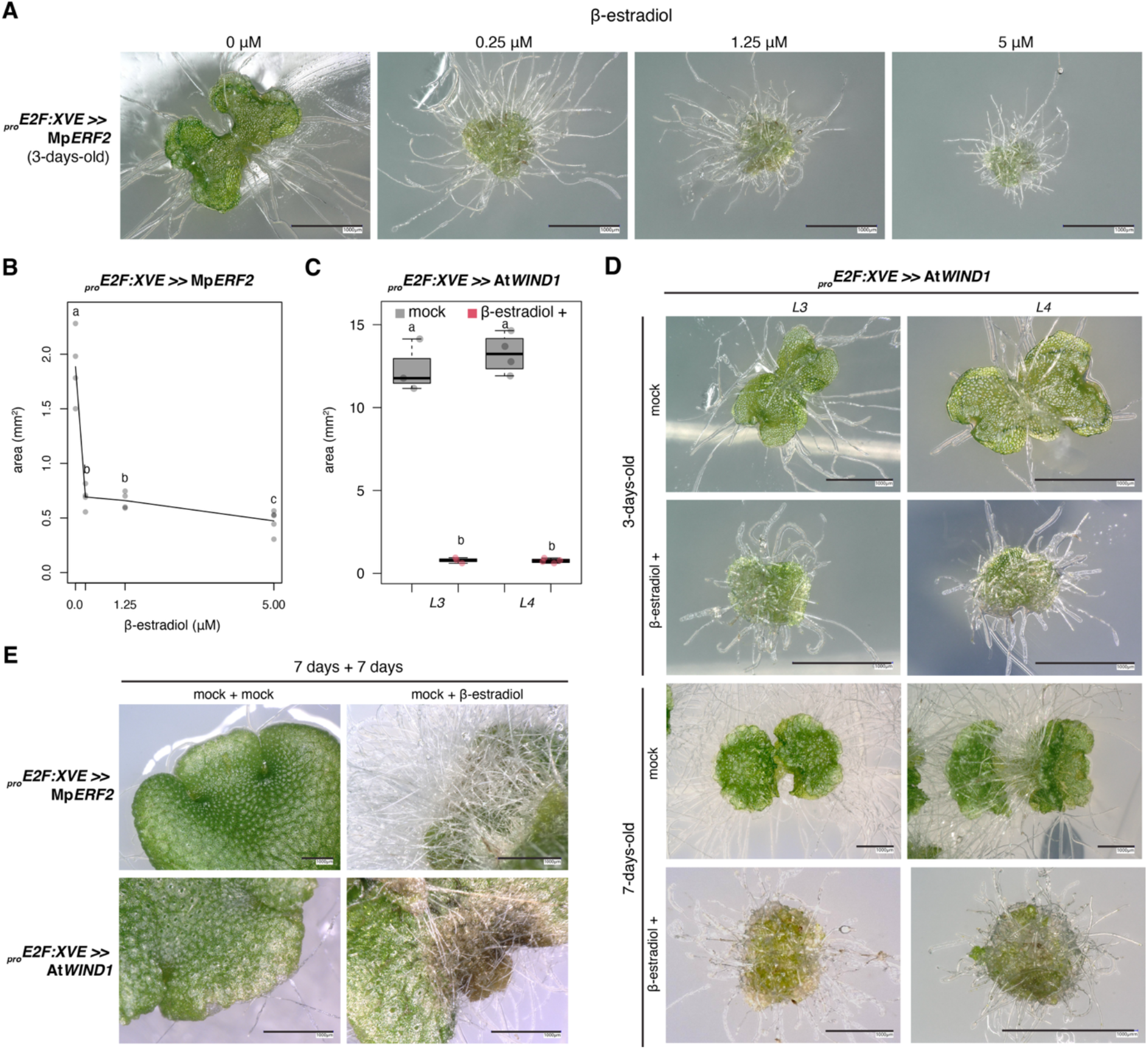
MpERF2 and AtWIND1 induce ectopic rhizoid formation. (A) Representative images of 3-day-old gemmalings expressing inducible MpERF2 (XVE >> MpERF2 L1) after induction with different concentrations of β-estradiol for 3 days as indicated. (B) Quantification of thallus area of the plants shown in panel A (n=4). (C) Quantification of thallus area in 7-day-old gemmalings expressing inducible AtWIND1 following induction (n=4). Two independent lines were used (L4 and L5). (D) Representative images of 3- and 7-day-old gemmalings expressing inducible AtWIND1 following induction. (E) Representative images following induction of MpERF2 and heterologous AtWIND1 expression at later developmental stages. Scale bars are indicated in each panel. Statistical significance was assessed using Tukey’s HSD test; groups with significantly different means are indicated by different letters (95% confidence level). Scale bars = 1000 μm.

To test whether the developmental activity of Group I ERFs is conserved, we heterologously expressed At*WIND1* in Marchantia. At*WIND1* also induced ectopic rhizoid-like outgrowths in 3- and 7-day-old gemmalings, although the phenotype was milder than that caused by MpERF2 (Figure 5C,D).

We next tested whether the activity of Mp*ERF2* and At*WIND1* was restricted to early gemmaling development. Plants were grown for 7 days without β-estradiol to allow the meristem to fully mature (38) before induction. Both MpERF2 and AtWIND1 induced ectopic rhizoid-like outgrowths on the dorsal thallus following induction (Figure 5E). This further strengthens the idea that Mp*ERF2* is sufficient to induce ectopic rhizoids across the developmental stages tested. The appearance of ectopic rhizoids across the differentiated dorsal thallus, suggest that MpERF2 activity is cell autonomous.

These results suggest that Group I ERFs retain a conserved capacity for cell fate determination that can be deployed toward different developmental outcomes. However, Group I ERFs have diversified extensively in Arabidopsis. MpERF2 is a putative orthologue of the 10 Arabidopsis Group I ERFs, which comprise several divergent clades. To further test this, we heterologously expressed AtERF54 and AtERF61, which represent two of the most divergent Group I ERF clades in Arabidopsis (Figure 1). In both cases, plants developed ectopic rhizoids and showed reduced thallus area (Supplemental Figure 6). However, unlike MpERF2- and AtWIND1-expressing plants, the apical notch persisted and was not fully converted into rhizoid-like tissue. Thus, AtERF54 and AtERF61 retain the capacity to induce rhizoid formation but the phenotypic outcomes differ from MpERF2 and AtWIND1, indicating some functional diversification within Group I ERFs.

### MpERF2 is important for normal rhizoid development

To assess the physiological function of MpERF2, we generated two independent knock-out lines in Marchantia using homologous-recombination-mediated gene targeting in the *Tak-1* background and two lines were obtained (Mp*erf2-2^ko^* and Mp*erf2-42^ko^*). The knock-out lines did not display conspicuous developmental defects compared to wild-type thallus. Yet, the area covered by rhizoids was significantly reduced, resulting in a lower rhizoid-to-thallus ratio area (Figure 6A,B). Loss of MpERF2 did not significantly affect regeneration compared with wild-type plants (Figure 6C-D). This corroborates that MpERF2 is dispensable for regeneration under the conditions tested but contributes to normal rhizoid differentiation, consistent with promoter analysis and gain-of-function phenotype.

**Figure 6.**
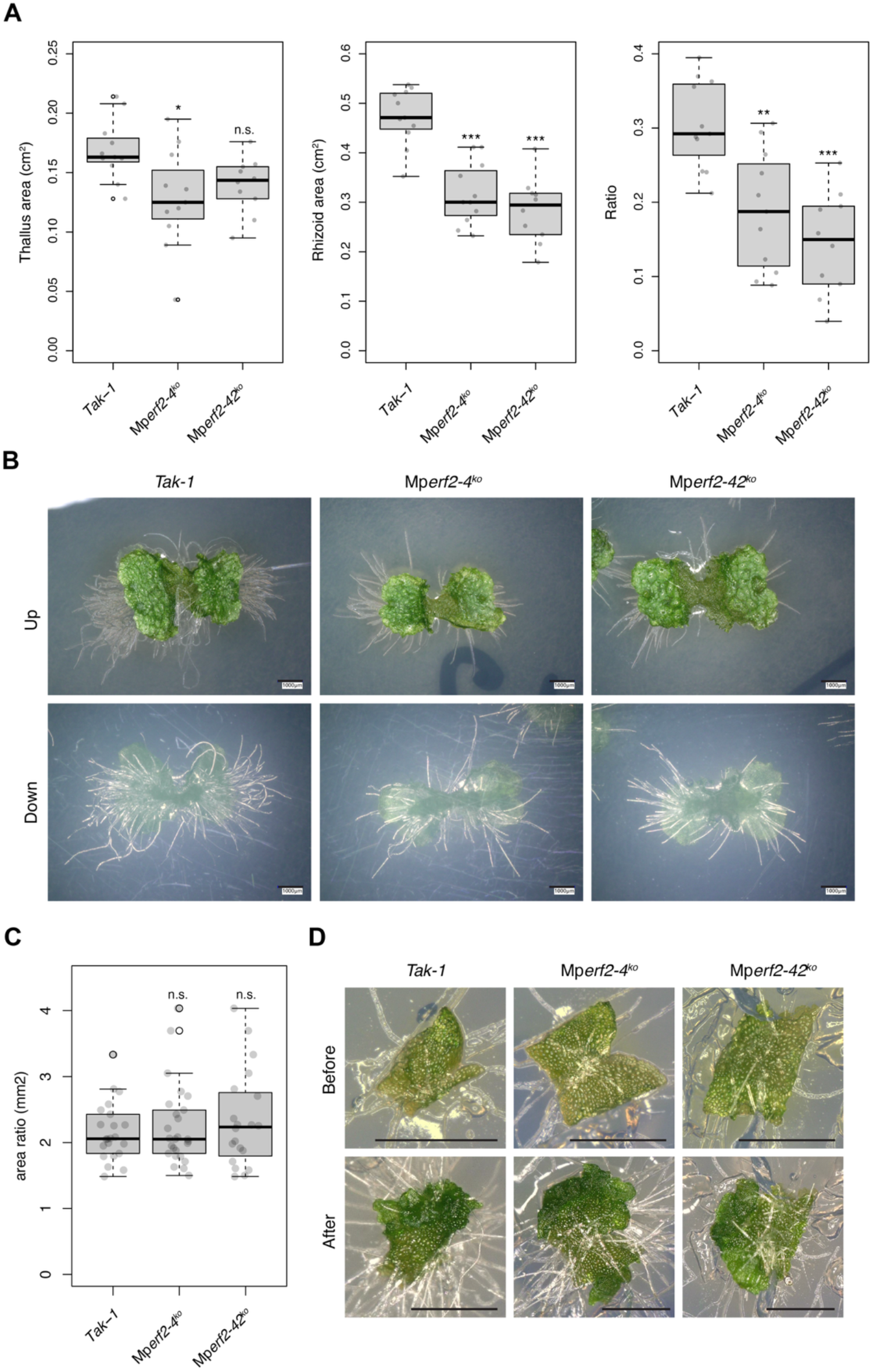
MpERF2 contributes to normal rhizoid development but is dispensable for regeneration. (A) Quantification of thallus area, rhizoid area, and rhizoid area-to-thallus area ratio in 7-day-old gemmalings of MpERF2 knockout lines and wild-type (Tak-1) plants. (B) Representative bright-field images of the plants quantified in (A). Scale bars = 1000 μm. (C) Regeneration of 3-day-old gemmalings following excision of the meristem in MpERF2 knockout lines and wild-type plants. Quantification of regenerated thallus area 4 days after excision. (D) Representative images before and 4 days after meristem excision corresponding to the plants shown in (C). Scale bars = 1000 μm. Asterisks indicate significant differences (p-value * < 0.05, ** < 0.01, *** < 0.001).

### Transcriptome profiling identifies MpERF2-responsive genes

To characterize the transcriptional response to Mp*ERF2* activation, we performed RNA-seq across the genotypes generated. Mp*ERF2* transcript levels were strongly increased after 4 hours induction and even more after 24 hours, confirming activation of the transgene (Figure 7A). Because the mock-treated inducible lines displayed numerous differentially expressed genes relative to wild type, likely reflecting basal transgene expression, subsequent differential-expression analyses used wild-type Cam-1 plants treated with β-estradiol as the reference control. Mp*ERF2* activation resulted in extensive transcriptional changes, with 2985 positive (FDR < 0.05, Fold-Change > 2) and 2662 negative (FDR < 0.05, Fold-Change < -2) differentially expressed genes. RNA-seq of Mp*erf2-42^ko^* knock-out line identified a further 721 differentially expressed genes relative to its corresponding wild-type control (Figure 7B).

**Figure 7.**
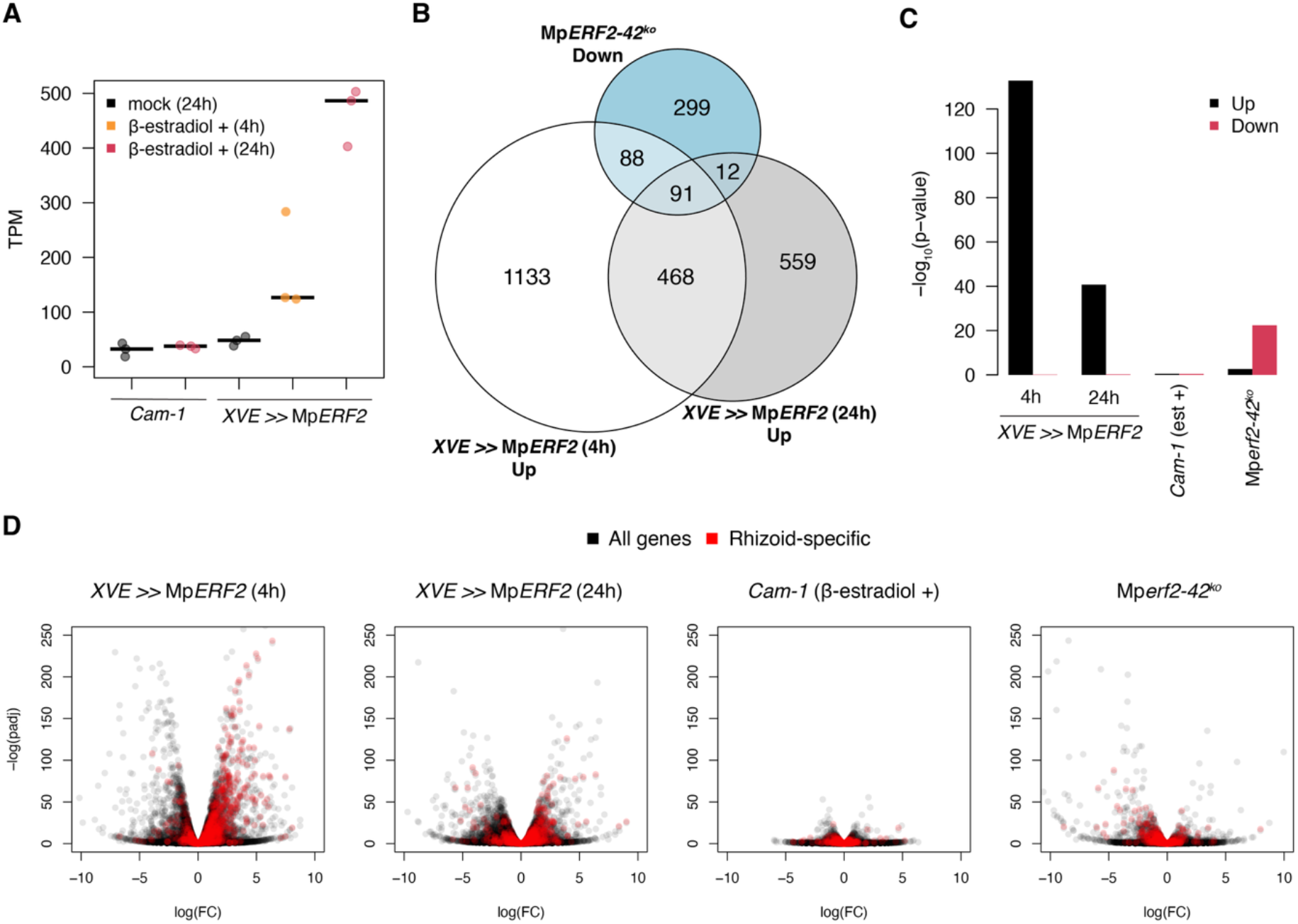
Transcriptomic changes associated with MpERF2 gain and loss of function. (A) MpERF2 expression following its induction in β-estradiol in wild type (Cam-1) and XVE >> MpERF2 (L1). (B) Overlap between differentially expressed genes identified in the MpERF2 gain- and loss-of-function transcriptomes. (C) Enrichment of rhizoid-specific genes among differentially expressed genes in MpERF2 gain- and loss-of-function plants. P-values correspond to Fisher-exact test. (D) Volcano plots showing differential gene expression in the MpERF2 gain- and loss-of-function transcriptomes. Rhizoid-specific genes are highlighted in red.

Relative to estradiol-treated wild type, MpERF2 activation was associated with broad transcriptome remodeling; notably, rhizoid-enriched genes were consistently upregulated (47), either at 4 hours or 24 hours (Figure 7C). Similarly, rhizoid-specific genes are significantly enriched in downregulated genes associated with the knock-out plants (Figure 7C). These genes include cell wall modifiers such as expansins (Mp*EXPANSIN3/4/5/12/17/25/31*) and aquaporins (MpPIP1/2, MpTIP2). Overall, this strongly supports the interpretation that the observed phenotypic changes are largely associated with a higher proportion of cells with rhizoids.

It was previously shown that the bHLH transcription factor Mp*RSL1* and its miRNA repressor Mp*FRH1* (48, 49) are important for rhizoid formation. Neither MpRSL1 nor its miRNA regulator MpFRH1 was differentially expressed (Supplemental Table S1), indicating that MpERF2 activity does not substantially alter the transcript abundance of MpRSL1 or MpFRH1. Thus, Mp*ERF2* relationship with this established regulatory module is unresolved, as it could act downstream or in parallel.

Among other transcription factors, Mp*ERF9* was upregulated following MpERF2 activation, whereas Mp*ERF21* was downregulated in both gain- and loss-of-function conditions. Both genes are related to Arabidopsis ERF115, suggesting that some connections of the transcriptional network associated with regeneration in Arabidopsis (8) are also observed in Marchantia.

## DISCUSSION

ERF transcription factors provide a striking example of how conserved transcriptional regulators can diversify to control distinct developmental programs. Beyond their canonical roles in stress signalling, ERFs have been recruited into diverse processes including cell differentiation. By systematically examining the ERF family in Marchantia, we were able to compare the phenotypic outcomes associated with representative from all groups. We found extensive functional diversification and identified Mp*ERF2* as a Group I ERF that promotes rhizoid development rather than regeneration. In Marchantia, this capacity to drive cellular fate is also shared by other members of this family, including Mp*ERF14/GMFR* in gemma formation (18), Mp*ERF13* in oil-body cell differentiation (17). Interestingly, each case is associated with specific cellular identities. Following the naming logic of Mp*ERF14/GMFR*, we propose to rename Mp*ERF2* as Mp*RHIZOFER* and Mp*ERF13* as Mp*ELAIOFER*.

Mp*ERF2* is the single group I member and the closest homologue of the Arabidopsis WIND clade. Our functional analysis contrasts with overexpression studies in *Arabidopsis*, where *WIND1* promotes dedifferentiation and enhances callus formation and somatic embryogenesis (13). Thus, Mp*ERF2* does not appear to be a major component of the regeneration program in Marchantia, unlike other members of this family, such as Mp*ERF20/LAXR* and Mp*ERF15*, which contribute to regeneration (10, 11). One important caveat is that in Arabidopsis, the direct link between wounding and regeneration is normally associated with group X ERFs (*ERF109/115*), while Mp*ERF15* belongs to group IX (Figure 1). As both groups contain ERFs associated with JA/ethylene signalling, this raises the possibility that related ERFs have been repeatedly recruited into wound-responsive regulatory networks (50).

More broadly, this work underscores the importance of bryophytes in reconstructing ancestral the functional evolution of transcription factors families and shed light on molecular mechanisms of functional innovations (51). These transcription factors share similar DNA binding preference and can be separated into two types of motifs: GCC (GCCGCC) or DRE (CCGAC). The first is associated with Groups VI-XI, and the second to Groups I-V (52). While ERFs involved in regeneration in both Marchantia and Arabidopsis, belong to the GCC group. The ERFs groups that we found to be associated with cell fate determination and reprogramming in Marchantia can be found in both.

The WIND clade of ERFs is a product of an expansion of Group I in flowering plants (Figure 1) and includes genes that are not associated with regeneration. This suggests that AtWIND1 might have been co-opted in the regeneration GRN to facilitate the process later in evolution. This evolutionary transition may have involved changes in downstream targets, but also in protein domains and regulatory interactions. The incorporation of the N1 motif into the WIND clade may have contributed to the evolution of its regulatory interactions in angiosperms. Notably, MpERF2 lacks this motif yet retains the capacity to induce rhizoid formation, indicating that this activity does not require N1. As shown recently, this motif is critical for WIND1’s capacity to interact with H3 acetylation/deacetylation machinery components (15). A similar type of molecular innovation could be associated with the lack of conservation of the group VII ERFs in oxygen sensing (32), suggesting that this mechanism might be a critical driver of neofunctionalization among ERFs.

Beyond differences in protein domains and regulatory interactions, the distinct developmental role of Mp*ERF2* occurs in the context of fundamentally different regenerative strategies in bryophytes and flowering plants. In Marchantia, regeneration proceeds without the typical parenchymatous callus stage and without exogenous hormonal treatments, commonly used to initiate callus formation and organogenesis in angiosperm tissue culture. These differences are accompanied by major differences in the downstream gene regulatory networks controlling meristem maintenance in the gametophyte compared with flowering plants (53, 54).

Root hairs and rhizoids are specialized tip-growing cells. They involve similar cell differentiation networks (49, 55), despite being present in the sporophyte and gametophyte, respectively. Heterologous expression of AtWIND1 in Marchantia also induced ectopic rhizoid formation. Interestingly, heterologous expression of AtERF54 and AtERF61 also display more rhizoids but the phenotypic outputs are distinct from those of MpERF2 and AtWIND1. However, no clear connection between Group I ERFs and root-hair formation has been established in Arabidopsis. These results suggest that Group I ERFs proteins retain a conserved capacity to engage a rhizoid-associated programme in the Marchantia heterologous context, despite their diversification. Whether this activity reflects an ancestral function of Group I ERFs or represents repeated recruitment of a conserved developmental capacity remains unresolved.

Another interesting example is the case of functional diversification of the group V ERFs, which are absent in Marchantia. Research in *P. patens* showed the group V ERFs (56) in the regulation of cuticle-associated programmes, similar to their orthologue in Arabidopsis *WAX INDUCER1/SHINE1* (*WIN1/SHN1*) proteins, which regulate cuticle-associated programs (57). Interestingly, other members of this group in *P. patens*, *STEM CELL-INDUCING FACTOR 1* (Pp*STEMIN1*) and its paralogues Pp*STEMIN2* and Pp*STEMIN3*, are critical for regeneration (58). This example illustrates how the recruitment of ERFs from different groups into the wound-induced regeneration pathway can also independently happen in bryophytes.

Several additional ERFs identified in our screen in Marchantia showed distinct developmental phenotypes and are also induced in regeneration, particularly Mp*ERF1*, Mp*ERF7* and Mp*ERF21*. This highlights additional candidates for the regulation of wound responses and regeneration in Marchantia. Defining their endogenous functions will help resolve the remaining nodes of the ERF regulatory networks that shaped land-plant evolution.

## ACKNOWLEDGMENTS

We thank Sakiko Ishida for generating the Mp*ERF2* knockout lines used in this study and Ryuichi Nishihama (Tokyo University of Science) and Yuki Hirakawa (Hiroshima University) for providing valuable feedback on the manuscript. This work was funded by Biotechnology and Biological Sciences Research Council (BB/F011458/1) for confocal microscopy. F.R. is a Leverhulme Early Career Fellow (ECF-2023-534) funded by the Leverhulme Trust and the Isaac Newton Trust (23.08(f)), I.B. is funded by the Herschel Smith Fund studentship.

## MATERIAL AND METHODS

### Phylogenetic analysis

*Amborella trichopoda* v1.0 (59), *Physcomitrium patens* v6.1 (60), and *Ceratopteris richardii* v2.1 (61) proteomes were obtained from Phytozome. The proteomes from *Marchantia polymorpha Tak* v6.1 (23), *Selaginella kraussiana* v2 (62), *Anthoceros agresti* Oxford (63), *Ginkgo biloba* v2 (64), *and Takakia lepidozioides* (65) from their respective sources. Hidden Markov Model matrix for the AP2 PFAMs (PF00847.25) was used as a query for searching using hmmsearch function from HMMER v3.4 (66) with default parameters. We used the public server at usegalaxy.org to analyse the data (67). Only sequences with p-value < 5E-7 were retained. As quality control, this approach recovered all ERFs annotated in Arabidopsis TAIR10 and Marchantia v6.1, except for At1g25470 and At5g67000, without false positives.

Using the hmmsearch output, Seqinr in R (68) package was used to retrieve the sequences from the proteomes. Finally, JalView (69) was used to align sequences with the MAFFT software (70) and trimming the alignment to include only columns with 50% coverage. Maximum-likelihood phylogenetic analysis was performed using IQtree 3 (71) with 1000 bootstrap replicates and default parameters as a preliminary tree. The best substitution model found was VT+R8 and subsequently used for the phylogenetic analysis. Using this tree, we excluded all AP2 and RAV sequences that did not belong to the ERF family, except for MpAPB (Mp8g11450) that was used as an outgroup to root the tree and repeated the alignment and phylogenetic analysis using the same VT+R8 model and 1000 bootstrap replicates. This final tree was rooted and visualised using iTOL v6 (72). A separate tree was generated using the same approach but with only *M. polymorpha*, *T. lepidozioides*, and *A. thaliana* proteomes as visualized in Figure 1. Both final trees are available at (Supplemental Figure S1).

The sequence DLNxxP was used to identify EAR motifs across the proteins and annotated in the phylogeny. The rest of the motifs were found using MEME (25) using default parameters.

### Plant material and growth conditions

*Marchantia polymorpha* subs*. rudelaris* accessions *Cam-1* (male) and *Cam-2* (female) were used for most of the experiments. Under normal conditions, plants were grown on solid 0.5× Gamborg B-5 basal medium (Phytotech #G398) at pH 5.8 with 1.2% (w/v) agar micropropagation grade (Phytotech #A296), under continuous LED light at 21 °C with light intensity of 150 *μ*mol/m^2^/s (Systion #SE-EGB). For spore production, plants were grown in Microbox micropropagation containers (SacO_2_) in long-day conditions (16 h light/8 h dark) under light supplemented with far-red light as described (73). _pro_MpERF2 in the Cam-1/2 background was described before (38).

### Plasmid construction

To generate new L0 parts, the CDS of Marchantia ERFs were extracted from *M. polymorpha* Tak-1 genome version 5.1 (5) and AtWIND1 from *A. thaliana* TAIR10 genome. Sequences were manually domesticated to remove internal BbsI, BsaI and SapI sites using synonymous mutations for the CDS. The sequences of synthetic L0 parts used in this work are available in Supplemental Table S2. L0 parts were synthesized by GENEWIZ following the standard syntax for plant synthetic biology with CDS12 overhangs and cloned into the plasmid pUAP1 (Addgene #63674) (74) by homology recombination. Other parts were derived from previous works (38, 73) as specified in Supplemental Table S2.

The acceptor pBy13 (75) was used to assemble the binary vectors using L0 corresponding to CDS12 and CTAG parts for β-estradiol inducible expression (XVE >>), as detailed in Supplemental Table S2. Cloning was performed using Type-IIS cloning as described previously for L3 plasmids (38).

### Plant transformation

*Agrobacterium tumefaciens* (GV3101) cells were transformed using the freeze-thaw method and used for plant transformation of *Cam-1/2* spores as described in Annese et al. (76) and selected for either 20 μg/mL hygromycin (Invitrogen #10687010) or 0.5μM chlorsulfuron (Thermo Scientific #17959385). At least 6 transgenic plants were screened for positive fluorescence and at least two independent lines were selected.

### Targeted gene knockout

To generate the targeting vectors, pJHY-TMp1 was used (77). The 3,547bp 5′ and the 3,587bp 3′ homology arms were amplified with the primer pairs, MpWIND1_5IF_L: ctaaggtagcgattaatCCGAAATTCTGCGTCTTCTC and MpWIND1_5IF_R: CCGGGCAAGCTTTTAAtCGTCTGCTGCTGCTGAATAA for the 5’ arm and MpWIND1_3IF_L: aacactagtggcgcgAGATTTCTCGTTGGGAGTGC and MpWIND1_3IF_R: TTATCCCTAGGCGCgCTTTGACTCGACCGACCTTC, and the PCR products were cloned into the *Asc*I and *Pac*I sites of pJHY-TMp1 using the In-Fusion HD cloning kit (Clontech), respectively. Introduction of the targeting construct into *M. polymorpha* and selection of knockout lines were performed as previously described (18, 78).

### Transcriptome analysis

For publicly available RNA-seqs, TPM values for sporeling germination and different developmental stages were extracted from Marpolbase Expression database (79) or FASTQ files were downloaded from SRA for regeneration (DRR330148-DRR330173), mapped in *M. polymorpha* Tak-1 genome v7.1 using HISAT2 (80), ht-seq and EdgeR using default parameters. Data was subsequently analysed with R to generate plots using custom scripts. Heatmap was generated using Marpolbase Expression database (79).

For new RNA-seq, XVE >> MpERF2 (L1) and wild-type (*Cam-1*) gemmalings were grown for 10 days in normal conditions and then 5mL of liquid Gamborg B5 liquid media supplemented with 5*μ*M β-estradiol (Merck #E8875) or without supplementation (mock) was added to the plates. Samples were collected either 4 hours or 24 hours, as indicated, before sample collection (n = 3). Similarly, samples from 10 days old gemmaling of Mp*erf2-42^ko^* and Mp*Tak-1* were collected (n =3). Total RNA was extracted using the Qiagen RNeasy Plant Mini Kit (Qiagen #74904) and enriched in PolyA selection for mRNA species. Library was prepared using the NEBNext Ultra II RNA kits (NEB) and module according to manufacturers instructions. Sequencing on a NovaSeq X plus (Illumina) at 2x150bp configuration by GENEWIZ Germany GmbH (Leipzig, Germany). Approximately 20 million paired-end reads per sample were obtained. FASTQ files were analysed as indicated before and differential expression analysis and plots were done using EdgeR glmLRT function (Bonferroni adjusted p-value < 0.01 and log2 fold-change > |1|) against their respective control.

### Plant phenotyping

Representative images of Marchantia plants were taken using a Keyence VHX-5000 digital microscope equipped with a 20x-200x Ultra-Small, High-Performance Zoom Lens (VH-Z20R/Z20T). For β-estradiol (Merck #E8875) treatments, a 50 mM stock was dissolved in DMSO and added to melted agar-Gamborg B-5 media at a final concentration of 5 μM as described in Ishida et al. (11). Plants were grown in microscopy contact plates (Simport Scientific #SIMPD210-17), and visualized after 3 or 10 days in normal growth conditions. Thallus area was quantified from pictures (3 or 7-days old gemmalings) with ImageJ software with at least 4-10 biological replicates. For regeneration, 3-days gemmalings were grown in contact plates, when meristems were excised and imaged after 4 days. Regenerating thallus area was quantified as described before (10) using pictures obtained from Keyence VHX-5000 digital microscope. BODIPY staining was performed as in Romani et al. (81) and imaged using a Leica epifluorescence DM600

### Laser scanning confocal microscopy

Confocal images of Marchantia were acquired on a Leica SP8X spectral confocal microscope upright system equipped with a 460 to 670 nm super continuum white light laser (80% laser power), two CW laser lines 405, and 442 nm, and 5 Channel Spectral Scanhead (four hybrid detectors and one photomultiplier). For slides, imaging was conducted using either a 10× air objective (HC PL APO 10×/0.40 CS2) or a 20× air objective (HC PL APO 20×/0.75 CS2). When observing fluorescent protein with overlapping emission spectra, sequential scanning mode was selected. Excitation laser wavelength and captured emitted fluorescence wavelength window were as follows: mVenus (514 nm, 527–552 nm), mScarlet-I (569 nm, 585–625 nm), and for chlorophyll autofluorescence (633, 687–739 nm).

For optical clearing, plants were fixed with 4% formaldehyde in PBS for 1 h under vacuum, washed with 1 mL PBS three times, then incubated with iTomei-D (#T3940, Tokyo Chemical Industry) overnight. Samples were washed again with 1 mL PBS and stained with SR2200 cell wall staining (Renaissance Chemicals, #2SCRI24) for 6 hours, washed in 1mL PBS and destained in PBS for 1 hour. Fixed, cleared and stained tissue was placed in 100 μL 70% iohexol (#I0903, Tokyo Chemical Industry) for 30 minutes, in which they were mounted inside of two stacked Gene Frames (Thermo Fisher Scientific). Samples were covered with a glass coverslip and imaged on a Leica SP8X confocal microscope as described before.

For _pro_MpERF2 time-courses, plants were grown under normal culture conditions in contact plates (Simport Scientific #SIMPD210-17), the lid was removed for imaging, and plants were returned to the growth chamber and imaged as described above.

### Statistics

To obtain plots and statistical analysis, data was processed using R version 4.4.1 software. For average and boxplots, the stats package was used with default parameters. In each boxplot, the central line indicates the median, and the box limits represent the first and third quartiles (interquartile range, IQR). Individual datapoints are indicated. For violinplots, the vioplot package was implemented also with default parameters. Statistical significance was calculated using ANOVA using the stats package Tukey ‘Honest Significant Difference’ method (alpha = 0.05) for significance levels testing.

---

**Supplemental Figure 1.**
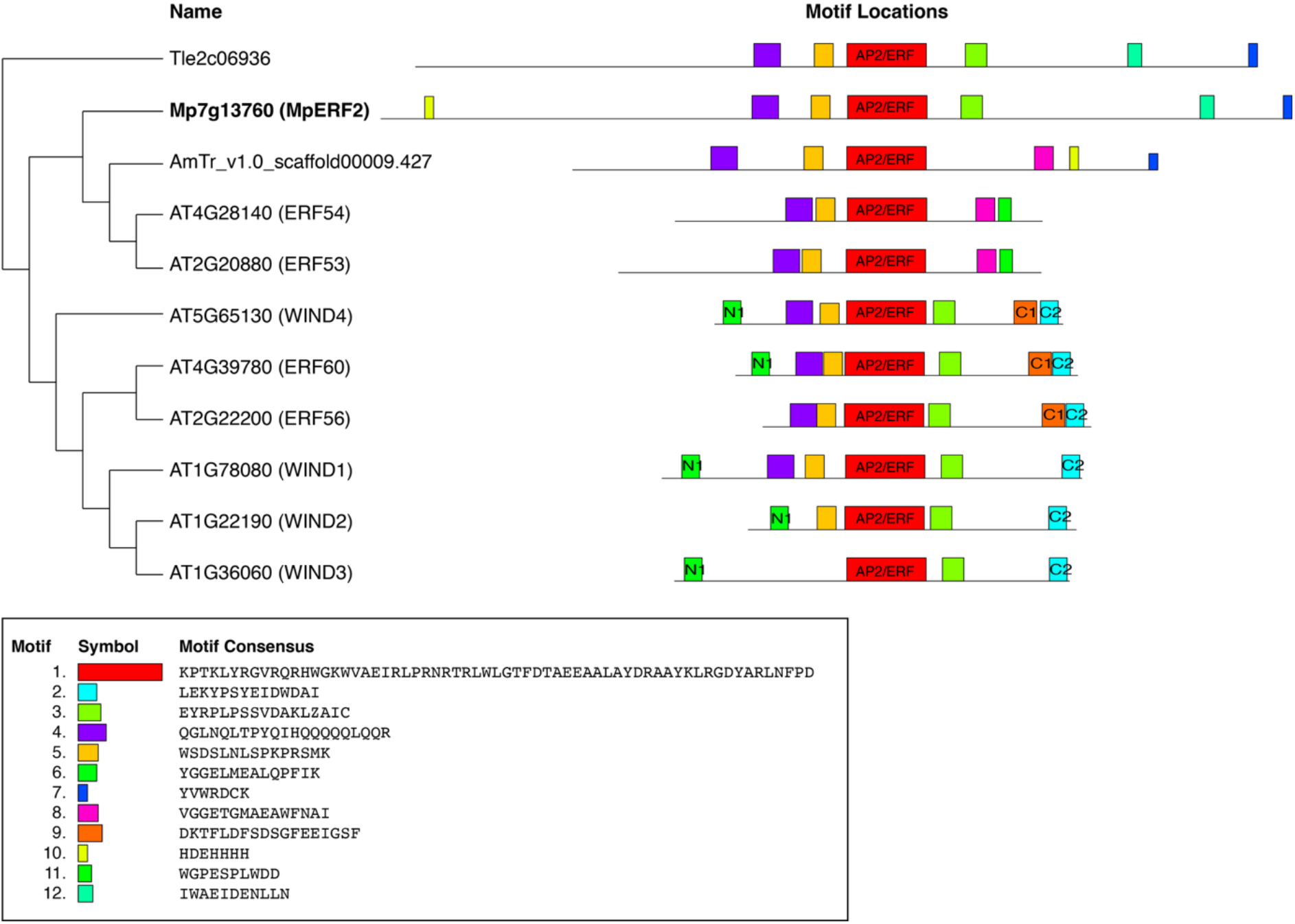
Conserved protein motifs in WIND-clade ERF transcription factors. MEME motif analysis of ERF proteins belonging to the WIND clade. Motifs are colored as shown in the legend. The light green corresponds to the conserved N1.

**Supplemental Figure 2.**
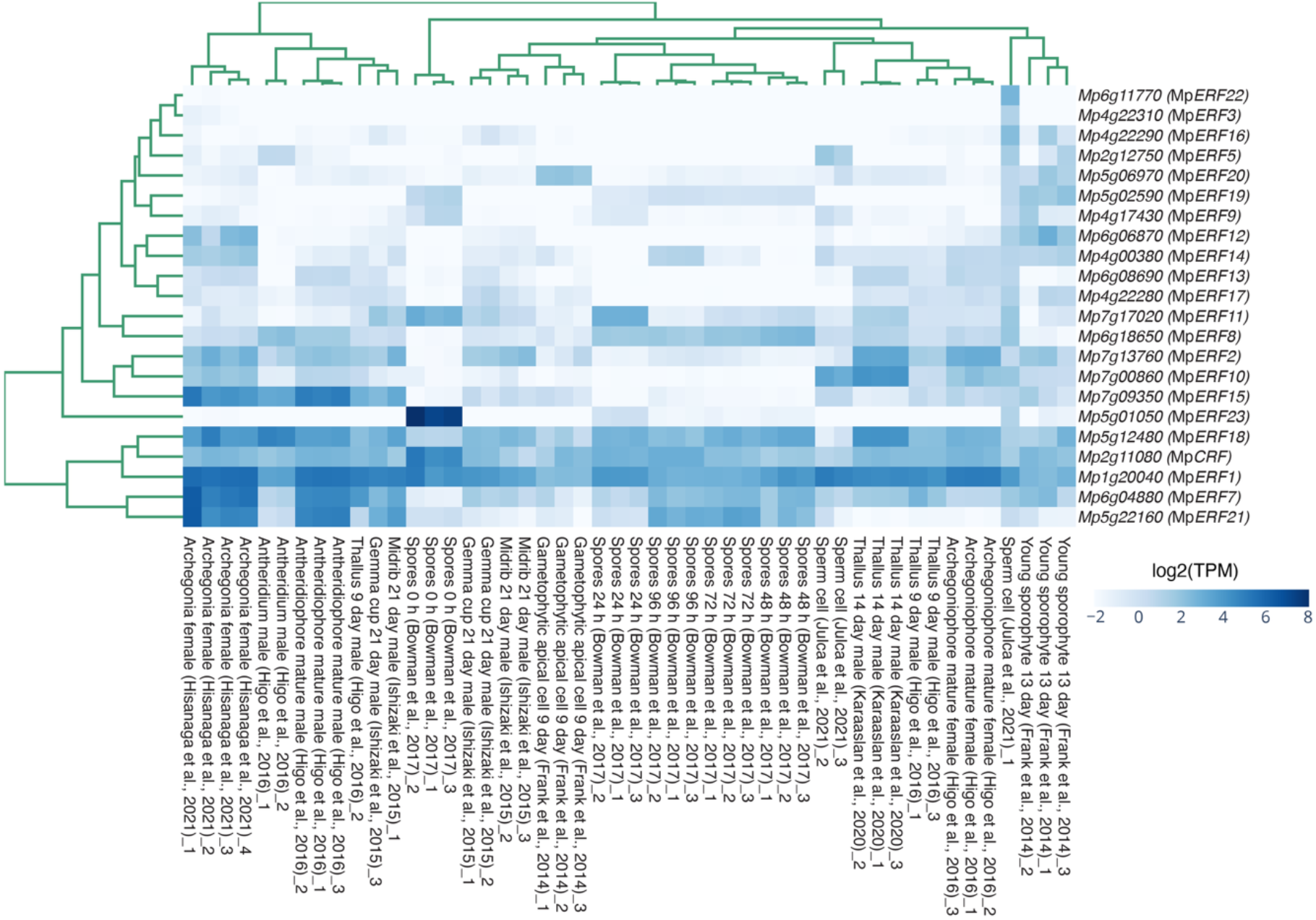
Expression profiles of ERF transcription factors across tissues and developmental stages of *Marchantia polymorpha*. The heatmap shows hierarchical clustering of ERF genes based on their expression profiles across different tissues and developmental stages, using data retrieved from the MarpolBase Expression (MBEX) database. Rows represent ERF genes, and columns represent individual tissues and developmental stages, as indicated by the labels below the heatmap. Rows and columns are reordered according to hierarchical clustering. The colour scale represents log₂-transformed transcripts per million (TPM) values, as indicated by the scale bar on the right.

**Supplemental Figure 3.**
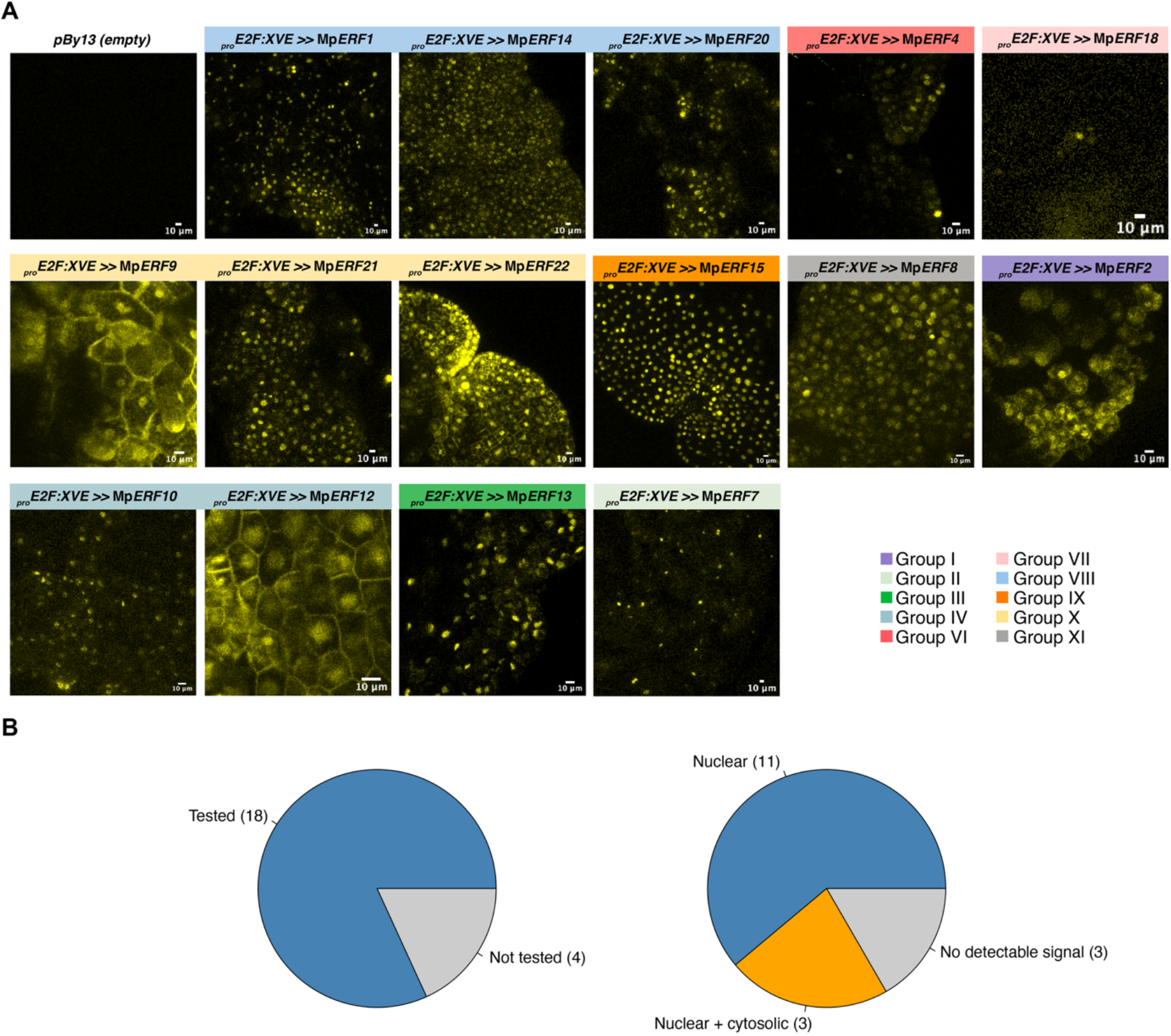
Marchantia ERF screening and subcellular localization. Representative confocal images maximum projection of Marchantia ERF inducible lines gemmalings in β-estradiol. pBy13 correspond to an empty vector control. Colour bars indicate ERFs groups according (see legend). (B) Pie charts summarising the screening and subcellular localization of Marchantia ERFs. Scale bars = 10 μm.

**Supplemental Figure 4.**
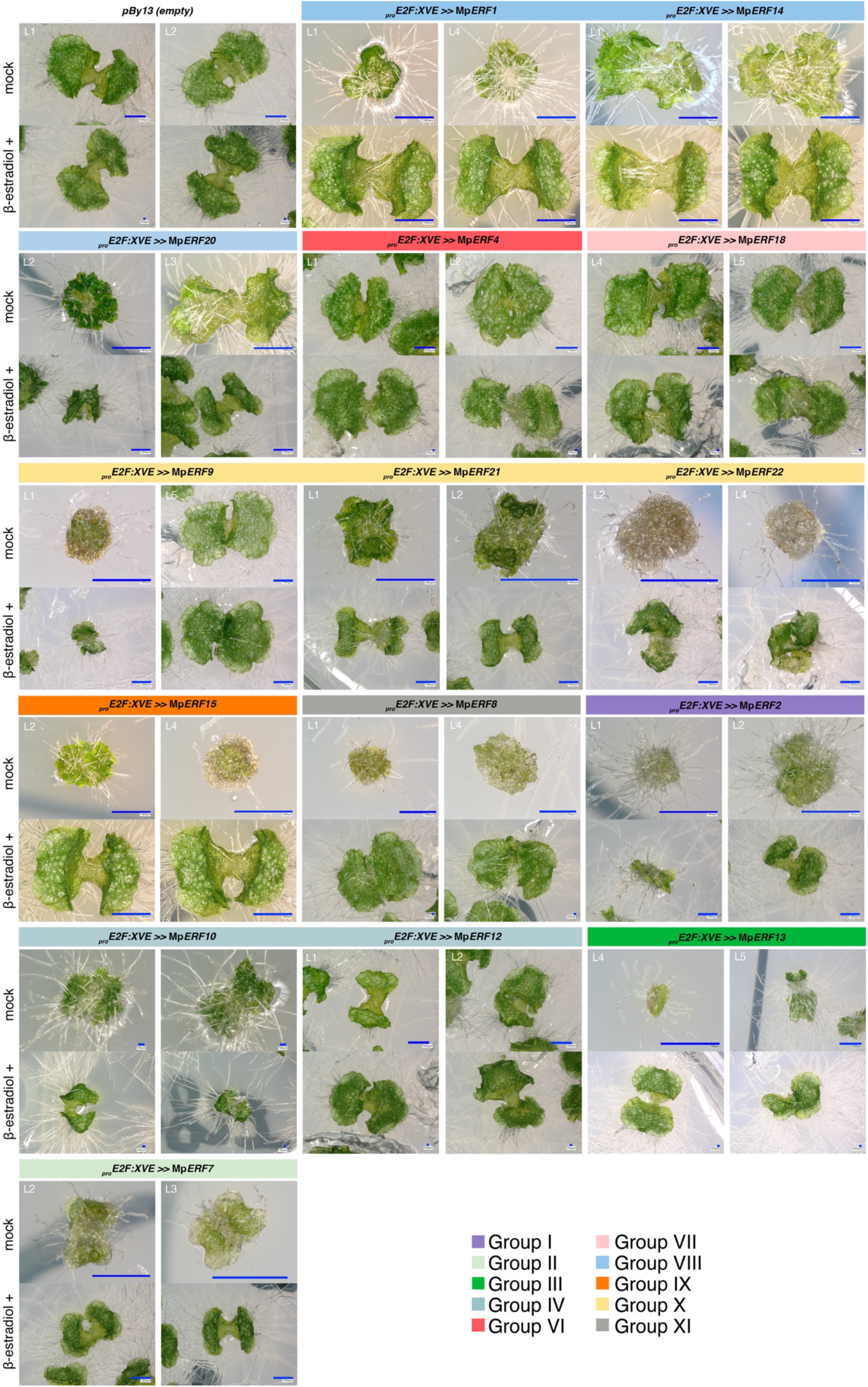
Systematic inducible overexpression of a subset of ERF transcription factors in Marchantia. Representative images of gemmalings of Marchantia ERF inducible lines after 7 days in Gamborg B5 media supplemented with 5 μM β-estradiol or without supplementation (mock). Two independent lines are shown for each genotype. pBy13 correspond to an empty vector control. Colour bars indicate ERFs groups according (see legend). Scale bars are indicated in each picture.

**Supplemental Figure 5.**
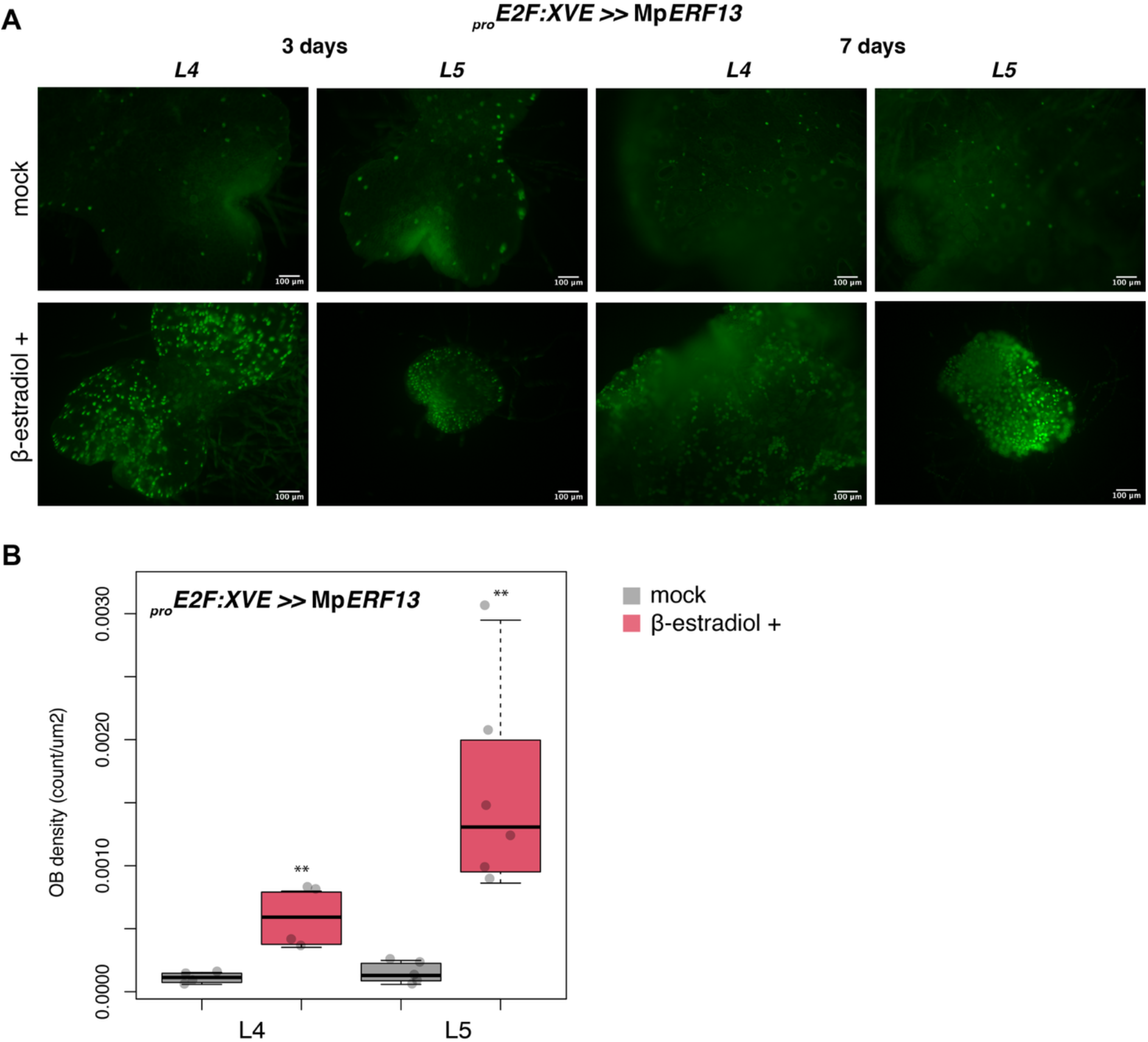
MpERF13 induces ectopic oil-body formation. (A) Representative images of MpE2F:XVE >> MpERF13 inducible lines stained with BODIPY to visualize oil bodies. Two independent lines are shown (L4 and L5). (B) Quantification of oil-body density (number of oil bodies per μm^2^) in each genotype and treatment (n=5). Asterisks indicate significant differences (p-value * < 0.05, ** < 0.01). Scale bars = 100 μm.

**Supplemental Figure 6.**
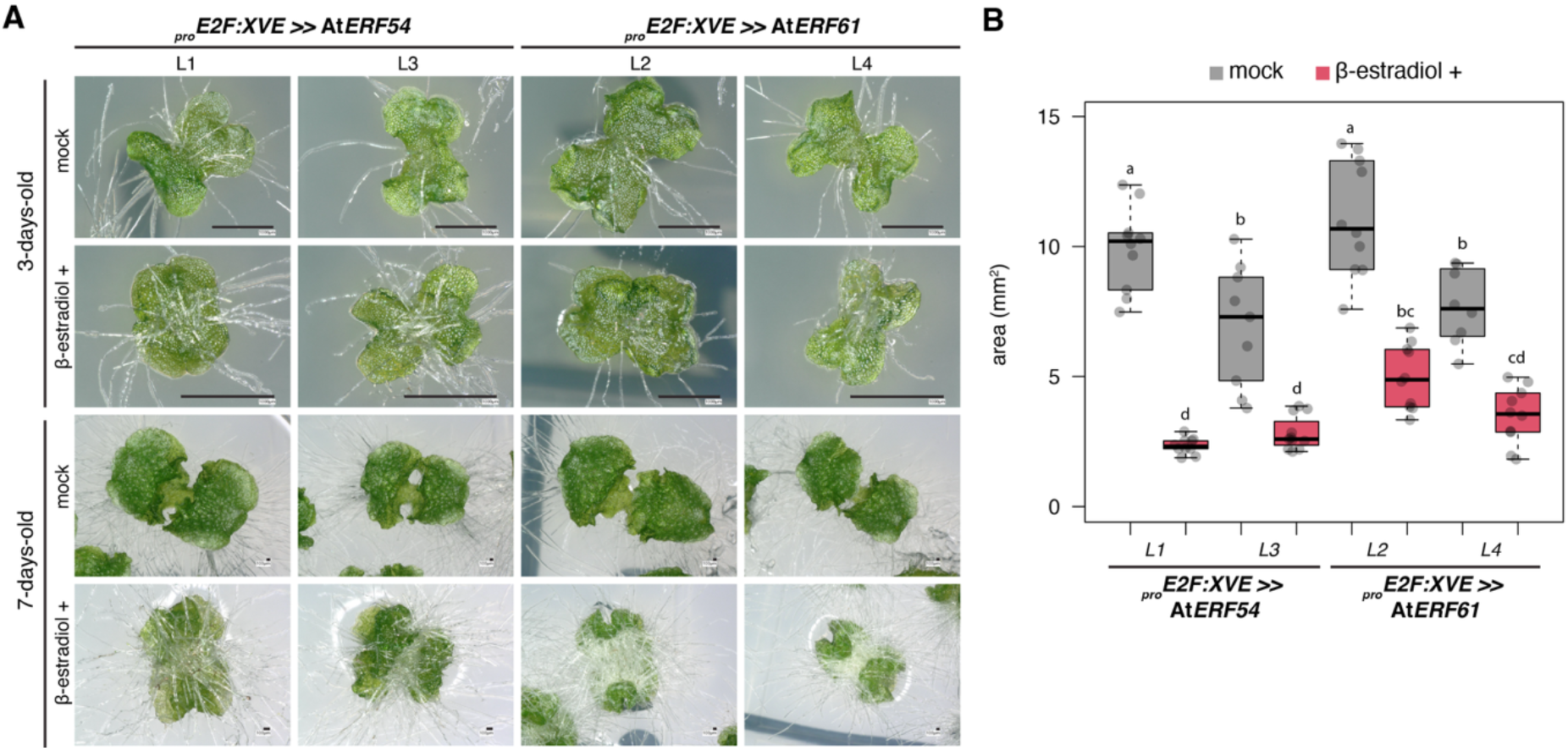
Heterologous expression of AtERF54 and AtERF61 in Marchantia. (A) Representative images of 3-day-old gemmalings expressing inducible AtERF54 or AtERF61 (XVE >> AtERF54 and XVE >> AtERF54 AtERF61) after induction with different concentrations of β-estradiol for 3 days as indicated. (B) Quantification of thallus area of the plants shown in 7-days-old gemmalings expressing inducible either AtERF54 or AtERF61 following induction as in panel A (n=10). Two independent lines were used per genotype. Scale bars are shown in each picture.

## REFERENCES

1. J. Jin et al., PlantTFDB 4.0: toward a central hub for transcription factors and regulatory interactions in plants. Nucleic Acids Res 45, D1040–D1045 (2017).

2. M. Ohme-Takagi, H. Shinshi, Ethylene-inducible DNA binding proteins that interact with an ethylene-responsive element. Plant Cell 7, 173–182 (1995).

3. F. Licausi, M. Ohme-Takagi, P. Perata, APETALA2/Ethylene Responsive Factor (AP2/ERF) transcription factors: mediators of stress responses and developmental programs. New Phytol 199, 639–649 (2013).

4. T. Nakano, K. Suzuki, T. Fujimura, H. Shinshi, Genome-wide analysis of the ERF gene family in Arabidopsis and rice. Plant Physiol 140, 411–432 (2006).

5. J. L. Bowman et al., Insights into Land Plant Evolution Garnered from the Marchantia polymorpha Genome. Cell 171, 287–304 e215 (2017).

6. J. Heyman, B. Canher, A. Bisht, F. Christiaens, L. De Veylder, Emerging role of the plant ERF transcription factors in coordinating wound defense responses and repair. J Cell Sci 131 (2018).

7. H. Morinaka et al., Transcriptome Dynamics of Epidermal Reprogramming during Direct Shoot Regeneration in Torenia fournieri. Plant Cell Physiol 62, 1335–1354 (2021).

8. M. Ikeuchi et al., A Gene Regulatory Network for Cellular Reprogramming in Plant Regeneration. Plant Cell Physiol 59, 765–777 (2018).

9. M. Ikeuchi, K. Sugimoto, A. Iwase, Plant callus: mechanisms of induction and repression. Plant Cell 25, 3159–3173 (2013).

10. Y. Liang et al., The wound-activated ERF15 transcription factor drives Marchantia polymorpha regeneration by activating an oxylipin biosynthesis feedback loop. Sci Adv 8, eabo7737 (2022).

11. S. Ishida et al., Diminished Auxin Signaling Triggers Cellular Reprogramming by Inducing a Regeneration Factor in the Liverwort Marchantia polymorpha. Plant Cell Physiol 63, 384–400 (2022).

12. R. Nishihama et al., Phytochrome-mediated regulation of cell division and growth during regeneration and sporeling development in the liverwort Marchantia polymorpha. J Plant Res 128, 407–421 (2015).

13. A. Iwase et al., The AP2/ERF transcription factor WIND1 controls cell dedifferentiation in Arabidopsis. Curr Biol 21, 508–514 (2011).

14. A. Iwase et al., WIND transcription factors orchestrate wound-induced callus formation, vascular reconnection and defense response in Arabidopsis. New Phytol 232, 734–752 (2021).

15. A. Iwase et al., WIND1 controls cell fate transition through coordinately integrating histone acetylation and deacetylation-mediated transcriptional reprogramming during somatic embryogenesis. Mol Plant 19, 1019–1035 (2026).

16. A. Iwase et al., WIND1 Promotes Shoot Regeneration through Transcriptional Activation of ENHANCER OF SHOOT REGENERATION1 in Arabidopsis. Plant Cell 29, 54–69 (2017).

17. T. Kanazawa et al., The liverwort oil body is formed by redirection of the secretory pathway. Nat Commun 11, 6152 (2020).

18. G. Takahashi et al., Initiation of asexual reproduction by the AP2/ERF gene GEMMIFER in Marchantia polymorpha. Curr Biol 36, 2413–2423 e2414 (2026).

19. G. Reboledo, A. Agorio, L. Vignale, A. Alvarez, I. Ponce De Leon, The moss-specific transcription factor PpERF24 positively modulates immunity against fungal pathogens in Physcomitrium patens. Front Plant Sci 13, 908682 (2022).

20. B. Causier, M. Ashworth, W. Guo, B. Davies, The TOPLESS interactome: a framework for gene repression in Arabidopsis. Plant Physiol 158, 423–438 (2012).

21. O. Lorenzo, R. Piqueras, J. J. Sanchez-Serrano, R. Solano, ETHYLENE RESPONSE FACTOR1 integrates signals from ethylene and jasmonate pathways in plant defense. Plant Cell 15, 165–178 (2003).

22. R. Solano, A. Stepanova, Q. Chao, J. R. Ecker, Nuclear events in ethylene signaling: a transcriptional cascade mediated by ETHYLENE-INSENSITIVE3 and ETHYLENE-RESPONSE-FACTOR1. Genes Dev 12, 3703–3714 (1998).

23. S. A. Montgomery et al., Chromatin Organization in Early Land Plants Reveals an Ancestral Association between H3K27me3, Transposons, and Constitutive Heterochromatin. Curr Biol 30, 573–588 e577 (2020).

24. S. Kagale, K. Rozwadowski, EAR motif-mediated transcriptional repression in plants: an underlying mechanism for epigenetic regulation of gene expression. Epigenetics 6, 141–146 (2011).

25. T. L. Bailey, C. Elkan, Fitting a mixture model by expectation maximization to discover motifs in biopolymers. Proc Int Conf Intell Syst Mol Biol 2, 28–36 (1994).

26. Z. Wang, S. J. Triezenberg, M. F. Thomashow, E. J. Stockinger, Multiple hydrophobic motifs in Arabidopsis CBF1 COOH-terminus provide functional redundancy in trans-activation. Plant Mol Biol 58, 543–559 (2005).

27. Y. Sakuma et al., Functional analysis of an Arabidopsis transcription factor, DREB2A, involved in drought-responsive gene expression. Plant Cell 18, 1292-1309 (2006).

28. Q. Liu et al., Two transcription factors, DREB1 and DREB2, with an EREBP/AP2 DNA binding domain separate two cellular signal transduction pathways in drought-and low-temperature-responsive gene expression, respectively, in Arabidopsis. Plant Cell 10, 1391–1406 (1998).

29. J. W. Cutcliffe, E. Hellmann, A. Heyl, A. M. Rashotte, CRFs form protein-protein interactions with each other and with members of the cytokinin signalling pathway in Arabidopsis via the CRF domain. J Exp Bot 62, 4995–5002 (2011).

30. A. M. Rashotte et al., A subset of Arabidopsis AP2 transcription factors mediates cytokinin responses in concert with a two-component pathway. Proc Natl Acad Sci U S A 103, 11081–11085 (2006).

31. F. Licausi et al., Oxygen sensing in plants is mediated by an N-end rule pathway for protein destabilization. Nature 479, 419–422 (2011).

32. L. Dalle Carbonare et al., ERFVIIs as transducers of oxygen-sensing in the evolution of land plant response to hypoxia. Mol Plant 18, 1072–1087 (2025).

33. S. Y. Fujimoto, M. Ohta, A. Usui, H. Shinshi, M. Ohme-Takagi, Arabidopsis ethylene-responsive element binding factors act as transcriptional activators or repressors of GCC box-mediated gene expression. Plant Cell 12, 393–404 (2000).

34. S. B. Tiwari et al., The EDLL motif: a potent plant transcriptional activation domain from AP2/ERF transcription factors. Plant J 70, 855–865 (2012).

35. A. Iwase et al., Arabidopsis WIND1 induces callus formation in rapeseed, tomato, and tobacco. Plant Signal Behav 8, e27432 (2013).

36. W. Liang et al., Dinor-12-oxo-phytodienoic acid conjugation with amino acids inhibits its phytohormone bioactivity in Marchantia polymorpha. Plant Physiol 197 (2024).

37. I. Monte et al., An Ancient COI1-Independent Function for Reactive Electrophilic Oxylipins in Thermotolerance. Curr Biol 30, 962–971 e963 (2020).

38. F. Romani et al., The landscape of transcription factor promoter activity during vegetative development in Marchantia. Plant Cell 36, 2140–2159 (2024).

39. P. Y. Huang, J. Catinot, L. Zimmerli, Ethylene response factors in Arabidopsis immunity. J Exp Bot 67, 1231–1241 (2016).

40. Y. Hata et al., snRNA-seq analysis of the moss Physcomitriumpatens identifies a conserved cytokinin-ESR module promoting pluripotent stem cell identity. Dev Cell 10.1016/j.devcel.2025.02.006 (2025).

41. H. Banno, Y. Ikeda, Q. W. Niu, N. H. Chua, Overexpression of Arabidopsis ESR1 induces initiation of shoot regeneration. Plant Cell 13, 2609–2618 (2001).

42. F. Y. Cao et al., Arabidopsis ETHYLENE RESPONSE FACTOR 8 (ERF8) has dual functions in ABA signaling and immunity. BMC Plant Biol 18, 211 (2018).

43. F. Romani et al., Liverwort oil bodies: diversity, biochemistry, and molecular cell biology of the earliest secretory structure of land plants. J Exp Bot 73, 4427–4439 (2022).

44. P. K. Agarwal, K. Gupta, S. Lopato, P. Agarwal, Dehydration responsive element binding transcription factors and their applications for the engineering of stress tolerance. J Exp Bot 68, 2135–2148 (2017).

45. H. Y. Chen et al., ORA47 (octadecanoid-responsive AP2/ERF-domain transcription factor 47) regulates jasmonic acid and abscisic acid biosynthesis and signaling through binding to a novel cis-element. New Phytol 211, 599–613 (2016).

46. M. J. Kim, D. Ruzicka, R. Shin, D. P. Schachtman, The Arabidopsis AP2/ERF transcription factor RAP2.11 modulates plant response to low-potassium conditions. Mol Plant 5, 1042–1057 (2012).

47. S. Kanno et al., Rhizoid-mediated phosphate uptake and internal transport in the non-vascular plant Marchantia polymorpha. New Phytol 250, 708–716 (2026).

48. A. Thamm, T. E. Saunders, L. Dolan, MpFEW RHIZOIDS1 miRNA-Mediated Lateral Inhibition Controls Rhizoid Cell Patterning in Marchantia polymorpha. Curr Biol 30, 1905–1915 e1904 (2020).

49. H. Proust et al., RSL Class I Genes Controlled the Development of Epidermal Structures in the Common Ancestor of Land Plants. Curr Biol 26, 93–99 (2016).

50. E. Flores-Sandoval, R. Nishihama, J. L. Bowman, Hormonal and genetic control of pluripotency in bryophyte model systems. Curr Opin Plant Biol 77, 102486 (2024).

51. F. Romani, J. E. Moreno, Molecular mechanisms involved in functional macroevolution of plant transcription factors. New Phytol 230, 1345–1353 (2021).

52. R. C. O’Malley et al., Cistrome and Epicistrome Features Shape the Regulatory DNA Landscape. Cell 165, 1280–1292 (2016).

53. E. Flores-Sandoval et al., The B-class auxin response factor MpARF2 is essential for meristem organization in free-living plant gametophytes. Curr Biol 10.1016/j.cub.2025.11.015 (2025).

54. I. R. Bonter, M., Delmans, M.; Romani, F.; Haseloff J., Marchantia stem cell maintenance and re-establishment are controlled by MpPIN1-mediated auxin transport and ARF signalling. BioRxiv 10.64898/2026.06.29.735252 (2026).

55. H. Breuninger et al., Diversification of a Transcription Factor Family Led to the Evolution of Antagonistically Acting Genetic Regulators of Root Hair Growth. Curr Biol 26, 1622–1628 (2016).

56. R. J. Kim, S. B. Lee, G. Pandey, M. C. Suh, Functional conservation of an AP2/ERF transcription factor in cuticle formation suggests an important role in the terrestrialization of early land plants. J Exp Bot 73, 7450–7466 (2022).

57. R. Kannangara et al., The transcription factor WIN1/SHN1 regulates Cutin biosynthesis in Arabidopsis thaliana. Plant Cell 19, 1278–1294 (2007).

58. M. Ishikawa et al., Physcomitrella STEMIN transcription factor induces stem cell formation with epigenetic reprogramming. Nat Plants 5, 681–690 (2019).

59. P. Amborella Genome, The Amborella genome and the evolution of flowering plants. Science 342, 1241089 (2013).

60. G. Bi et al., Near telomere-to-telomere genome of the model plant Physcomitrium patens. Nat Plants 10, 327–343 (2024).

61. D. B. Marchant et al., Dynamic genome evolution in a model fern. Nat Plants 8, 1038–1051 (2022).

62. W. Liu et al., Genome and transcriptome of Selaginella kraussiana reveal evolution of root apical meristems in vascular plants. Curr Biol 33, 4085–4097 e4085 (2023).

63. F. W. Li et al., Anthoceros genomes illuminate the origin of land plants and the unique biology of hornworts. Nat Plants 6, 259–272 (2020).

64. H. Liu et al., The nearly complete genome of Ginkgo biloba illuminates gymnosperm evolution. Nat Plants 7, 748–756 (2021).

65. R. Hu et al., Adaptive evolution of the enigmatic Takakia now facing climate change in Tibet. Cell 186, 3558–3576 e3517 (2023).

66. R. D. Finn, J. Clements, S. R. Eddy, HMMER web server: interactive sequence similarity searching. Nucleic Acids Res 39, W29–37 (2011).

67. C. Galaxy, Galaxy for accessible, reproducible, and collaborative data analyses: 2026 update. Nucleic Acids Res 54, W105–W116 (2026).

68. D. Charif, J. R. Lobry, "SeqinR 1.0-2: A Contributed Package to the R Project for Statistical Computing Devoted to Biological Sequences Retrieval and Analysis" in Structural Approaches to Sequence Evolution: Molecules, Networks, Populations, U. Bastolla, M. Porto, H. E. Roman, M. Vendruscolo, Eds. (Springer Berlin Heidelberg, Berlin, Heidelberg, 2007), 10.1007/978-3-540-35306-5_10, pp. 207-232.

69. J. B. Procter et al., Alignment of Biological Sequences with Jalview. Methods Mol Biol 2231, 203–224 (2021).

70. K. Katoh, K. Misawa, K. Kuma, T. Miyata, MAFFT: a novel method for rapid multiple sequence alignment based on fast Fourier transform. Nucleic Acids Res 30, 3059–3066 (2002).

71. B. Q. Minh et al., IQ-TREE 2: New Models and Efficient Methods for Phylogenetic Inference in the Genomic Era. Mol Biol Evol 37, 1530–1534 (2020).

72. I. Letunic, P. Bork, Interactive Tree of Life (iTOL) v6: recent updates to the phylogenetic tree display and annotation tool. Nucleic Acids Res 52, W78–W82 (2024).

73. S. Sauret-Gueto et al., Systematic Tools for Reprogramming Plant Gene Expression in a Simple Model, Marchantia polymorpha. ACS Synth Biol 9, 864–882 (2020).

74. N. J. Patron et al., Standards for plant synthetic biology: a common syntax for exchange of DNA parts. New Phytol 208, 13–19 (2015).

75. F. Romani et al., A simple cell-cycle control system in Marchantia polymorpha provides a framework for understanding plant cell proliferation. Plant Cell 10.1093/plcell/koag103 (2026).

76. D. Annese et al., Semi-automated workflow for high-throughput Agrobacterium-mediated plant transformation. Plant J 122, e70118 (2025).

77. K. Ishizaki, Y. Johzuka-Hisatomi, S. Ishida, S. Iida, T. Kohchi, Homologous recombination-mediated gene targeting in the liverwort Marchantia polymorpha L. Sci Rep 3, 1532 (2013).

78. K. Ishizaki, S. Chiyoda, K. T. Yamato, T. Kohchi, Agrobacterium-mediated transformation of the haploid liverwort Marchantia polymorpha L., an emerging model for plant biology. Plant Cell Physiol 49, 1084–1091 (2008).

79. S. Kawamura et al., MarpolBase Expression: A Web-Based, Comprehensive Platform for Visualization and Analysis of Transcriptomes in the Liverwort Marchantia polymorpha. Plant Cell Physiol 63, 1745–1755 (2022).

80. D. Kim, J. M. Paggi, C. Park, C. Bennett, S. L. Salzberg, Graph-based genome alignment and genotyping with HISAT2 and HISAT-genotype. Nat Biotechnol 37, 907–915 (2019).

81. F. Romani et al., Oil Body Formation in Marchantia polymorpha Is Controlled by MpC1HDZ and Serves as a Defense against Arthropod Herbivores. Curr Biol 30, 2815–2828 e2818 (2020).

