## Supplementary figures and images for "Functional diversification of ERF transcription factors in land plants"

### Supplemental File 1

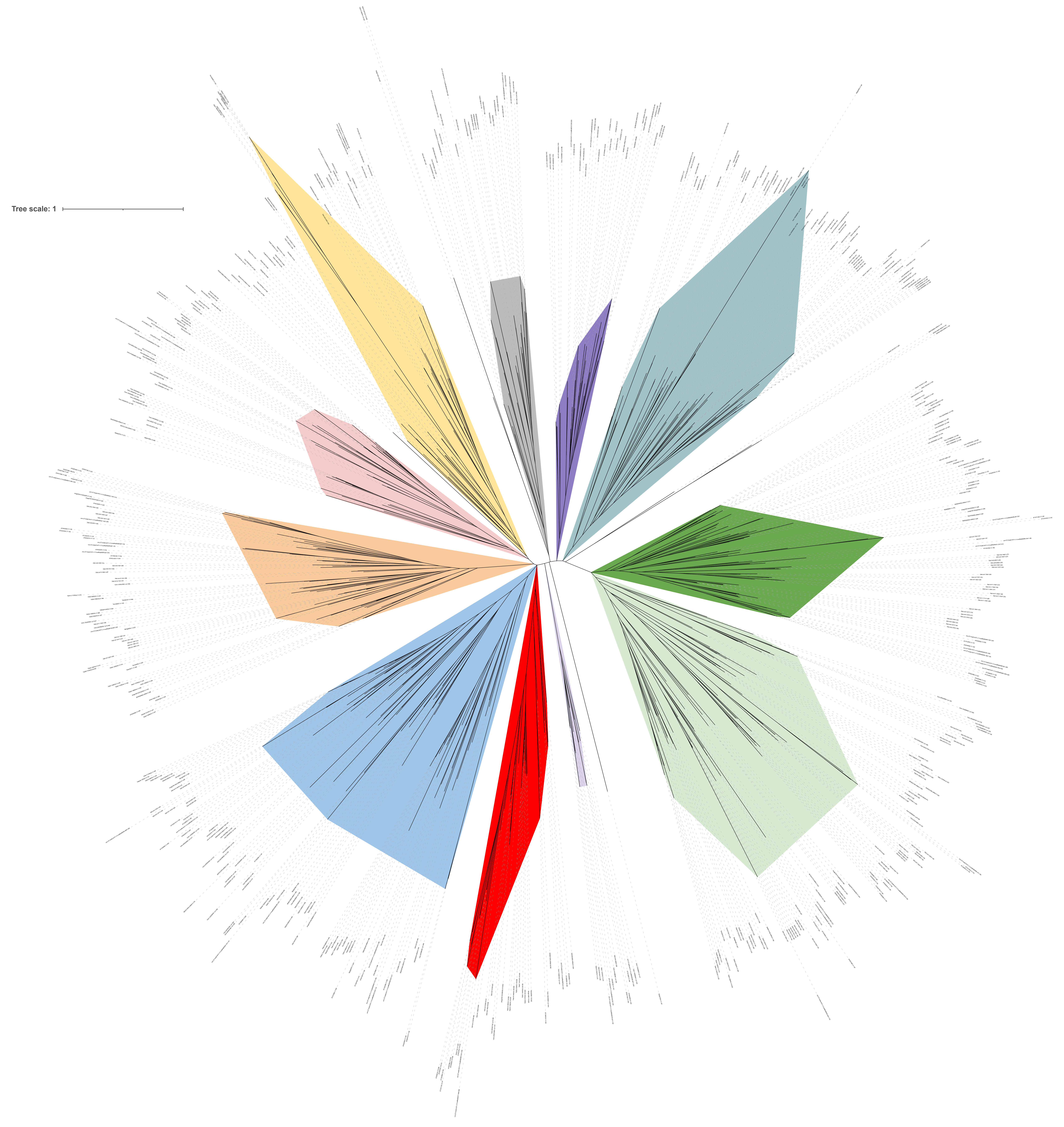
